# Handling Multiplicity in Neuroimaging through Bayesian Lenses with Multilevel Modeling

**DOI:** 10.1101/238998

**Authors:** Gang Chen, Yaqiong Xiao, Paul A. Taylor, Justin K. Rajendra, Tracy Riggins, Fengji Geng, Elizabeth Redcay, Robert W. Cox

## Abstract

Here we address the current issues of inefficiency and over-penalization in the massively univariate approach followed by the correction for multiple testing, and propose a more efficient model that pools and shares information among brain regions. Using Bayesian multilevel (BML) modeling, we control two types of error that are more relevant than the conventional false positive rate (FPR): incorrect sign (type S) and incorrect magnitude (type M). BML also aims to achieve two goals: 1) improving modeling efficiency by having one integrative model and thereby dissolving the multiple testing issue, and 2) turning the focus of conventional null hypothesis significant testing (NHST) on FPR into quality control by calibrating type S errors while maintaining a reasonable level of inference efficiency The performance and validity of this approach are demonstrated through an application at the region of interest (ROI) level, with all the regions on an equal footing: unlike the current approaches under NHST, small regions are not disadvantaged simply because of their physical size. In addition, compared to the massively univariate approach, BML may simultaneously achieve increased spatial specificity and inference efficiency, and promote results reporting in totality and transparency. The benefits of BML are illustrated in performance and quality checking using an experimental dataset. The methodology also avoids the current practice of sharp and arbitrary thresholding in the *p*-value funnel to which the multidimensional data are reduced. The BML approach with its auxiliary tools is available as part of the AFNI suite for general use.

## Introduction

The typical neuroimaging data analysis at the whole brain level starts with a preprocessing pipeline, and then the preprocessed data are fed into a voxel-wise time series regression model for each subject. An effect estimate is then obtained at each voxel as a regression coefficient that is, for example, associated with a task/condition or a contrast between two effects or a linear combination among multiple effects. Such effect estimates from individual subjects are next incorporated into a population model for generalization, which can be parametric (e.g., Student’s t-test, AN(C)OVA, univariate or multivariate GLM, linear mixed-effects (LME) or Bayesian model) or nonparametric (e.g., permutations, bootstrapping, rank-based testing). In either case, this generally involves one or more statistical tests at each spatial element separately.

### Issues with controlling false positives

As in many scientific fields, the typical neuroimaging analysis has traditionally been conducted under the framework of null hypothesis significance testing (NHST). As a consequence, a big challenge when presenting the population results is to properly handle the multiplicity issue resulting from the tens of thousands of simultaneous inferences, but this undertaking is met with various subtleties and pitfalls due to the complexities involved: the number of voxels in the brain (or a restricting mask) or the number of nodes on surface, spatial heterogeneity, violation of distributional assumptions, etc. The focus of the present work will be on developing an efficient approach from Bayesian perspective to address the multiplicity issue as well as some of the pitfalls associated with NHST (Appendix A). We first describe the multiplicity issue and how it directly results from the NHST paradigm and inefficient modeling, and then translate many of the standard analysis features to the proposed Bayesian framework.

Following the conventional statistical procedure, the assessment for a BOLD effect is put forward through a null hypothesis *H*_0_ as the devil’s advocate; for example, an *H*_0_ can be formulated as having no activation at a brain region under, for example, the easy condition, or as having no activation difference between the easy and difficult conditions. It is under such a null setting that statistics such as Student’s t- or F-statistic are constructed, so that a standard distribution can be utilized to compute a conditional probability that is the chance of obtaining a result equal to, or more extreme than, the current outcome if *H*_0_ is imagined as the ground truth. The rationale is that if this conditional probability is small enough, one may feel comfortable in rejecting the straw man *H*_0_ and in accepting the alternative at a tolerable risk.

While NHST may be a reasonable formulation under some scenarios, there is a long history of arguments that emphasize the mechanical and interpretational problems with NHST (e.g., Cohen, 2014; Gelman, 2016) that might have perpetuated the reproducibility crisis across many disciplines (Loken and Gelman, 2017). Within neuroimaging specifically, there are strong indications that a large portion of task-related BOLD activations are usually unidentified at the individual subject level due to the lack of power (Gonzalez-Castillo et al., 2012). The detection failure, or false negative rate, at the population level would probably be at least as large. Therefore, it is likely far-fetched to claim that no activation or no activation difference exists anywhere in the whole brain, except for the regions of white matter and cerebrospinal fluid. In other words, the global null hypothesis in enuroimaging studies is virtually never true. The situation with resting-state data analysis is likely worse than with task-related data, as the same level of noise is more impactful on seed-based correlation analysis due to the lack of objective reference effect. Since no ground truth is readily available, dichotomous inferences under NHST as to whether an effect exists in a brain region are intrinsically problematic, and it is practically implausible to truly believe the validity of H0 as a building block when constructing a model.

Achieving statistical significance has been widely used as the standard screening criterion in scientific results reporting as well as in the publication reviewing process. The difficulty in passing a commonly accepted threshold with noisy data may elicit a hidden misconception: A statistical result that survives the strict screening with a small sample size seems to gain an extra layer of strong evidence, as evidenced by phrases in the literature such as “despite the small sample size” or “despite limited statistical power.” However, when the statistical power is low, the inference risks can be perilous, as demonstrated with two different types of error as illustrated in Appendix B from the conventional type I and type II errors: incorrect sign (type S) and incorrect magnitude (type M). The conventional concept of FPR controllability is not a well-balanced choice under all circumstances or combinations of effect and noise magnitudes. We consider a type S error to be more severe than a type M error, and thus we aim to control the former while at the same time reducing the latter as much as possible, parallel to the similarly lopsided strategy of strictly controlling type I errors at a tolerable level under NHST while minimizing type II errors.

### Issues with handling multiplicity

In statistics, multiplicity is more often referred to as multiple comparisons or multiple testing problem when more than one statistical inference is made simultaneously (Appendix C). The challenges of dealing with multiple testing at the voxel or node level have been recognized within the neuroimaging community almost as long as the history of FMRI. Substantial efforts have been devoted to ensuring that the actual type I error (or FPR) matches its nominal requested level under NHST. Due to the presence of spatial non-independence of noise, the classical approach to countering multiple testing through Bonferroni correction in general is highly conservative when applied to neuroimaging, so the typical correction efforts have been channeled into two main categories, 1) controlling for FWE, so that the overall FPR at the cluster or whole brain level is approximately at the nominal value, and 2) controlling for false discovery rate (FDR), which harnesses the expected proportion of identified items or discoveries that are incorrectly labeled (Benjamini and Hochberg, 1995). FDR can be used to handle a needle-in-haystack problem where a small number of effects existing among a sea of zero effects in, for example, bioinformatics. However, FDR is usually quite conservative for typical neuroimaging data and thus is not widely adopted. Therefore, we do not discuss it hereafter in the current context.

Typical FWE correction methods for multiple testing include Monte Carlo simulations (Forman et al., 1995), random field theory (Worsley et al., 1992), and permutation testing (Nichols and Holmes, 2001; Smith and Nichols, 2009). Regardless of the specific FWE correction methodology, the common theme is to use the spatial extent, either explicitly or implicitly, as a pivotal factor. One recent study suggested that the nonparametric methods seem to have achieved a more uniformly accurate controllability for FWE than their parametric counterparts (Eklund et al., 2016), even though parametric methods may exhibit more flexibility in modeling capability (and some parametric methods can show reasonable FPR controllability; Cox et al., 2017). Because of this recent close examination (Eklund et al., 2016) on the practical difficulties of parametric approaches herein controlling FWE, there is currently an implied rule of thumb (Yeung, 2018) that demands any parametric correction be based on a voxel-wise p-value threshold at 0.001 or less. Such a narrow modeling choice with a harsh cutoff could be highly limiting, depending on several parameters such as trial duration (event-related versus block design), and would definitely make small regions even more difficult to pass through the NHST filtering system. In other words, the leverage on spatial extent with a Procrustean criterion undoubtedly incurs a collateral damage: small regions (e.g., amygdala) or subregions within a brain area are inherently placed in a disadvantageous position even if small regions have similar signal strength as larger ones; that is, to be able to surpass the same threshold bar, small regions would have to reach a much higher signal strength to survive a uniform criterion at the cluster threshold or whole brain level.

The concept of using contiguous spatial extent as a leveraging mechanism to control for multiplicity can be problematic from another perspective. For example, suppose that two anatomically separate regions are spatially distant and the statistical evidence (as well as signal strength) for each of their effects is not strong enough to pass the cluster correction threshold individually. However, if another two anatomically regions that have exactly the same statistical evidence (as well as signal strength) are adjacent, their spatial contiguity could elevate their combined volume to the survival of correction for FWE. Trade-offs are inherently involved in these final interpretations. One may argue that the sacrifice in statistical power under NHST is worth the cost in achieving the overall controllability of type I error, but it may be unnecessarily over-penalizing to stick to such an inflexible criterion rather than utilizing the neurological context or prior knowledge, as discussed below.

To summarize the debate surrounding cluster inferences, multiplicity is directly associated with the concept of false positives or type I errors under NHST, and the typical control for FWE at a preset threshold (e.g., 0.05, the implicitly accepted tolerance level in the field) is usually considered a safeguard for reproducibility. Imposing a threshold on cluster size (perhaps combined with signal strength) to protect against the overall FPR has the undesirable trade-off cost of inflating false negative rates or type II errors, which can greatly affect individual result interpretations as well as reproducibility across studies. In general, several multiplicity-related challenges in neuroimaging appear to be tied closely to the fundamental mechanisms of NHST approaches introduced to counterbalance between two counterfactual errors (type I and type II), which are the cornerstones of NHST. Therefore, we put forward a list of potential problems with NHST in Appendix A.

### Structure of the work

In light of the aforementioned backdrop, we believe that the current modeling approach is inefficient. First, we question the appropriateness of the severe penalty currently levied to the voxel- or node-wise data analysis. In addition, we endorse the ongoing statistical debate surrounding the ritualization of NHST and its dichotomous approach to results reporting and in the review process, and aim to raise the awareness of the issues embedded within NHST (Loken and Gelman, 2017) in the neuroimaging community. In addition, with the intention of addressing some of the issues discussed above, we view multiple testing as a problem of inefficient modeling induced by the conventional massively univariate methodology. Specifically, the univariate approach starts, in the same vein as a null hypothesis setting, with a pretense of spatial independence, and proceeds with many isolated or segmented models. To avoid the severe penalty of Bonferroni correction while recovering from or compensating for the false presumption of spatial independence, the current practices deal with multiple testing by discounting the number of models due to spatial relatedness. However, the collateral damages incurred by this to-and-fro process are unavoidably the loss of modeling efficiency and the penalty for detection power under NHST.

Here, we propose a more efficient approach through BML that could be used to confirm, complement or replace the standard NHST method. As a first step, we adopt a group analysis strategy under the Bayesian framework through multilevel modeling on an ensemble of ROIs and use this to resolve two of the four multiplicity issues above: multiple testing and double sidedness (Appendix C). Those ROIs are determined independently from the current data at hand, and they can be selected through various methods such as previous studies, an anatomical or functional atlas, or parcellation of an independent dataset in a given study; the regions could be defined through masking, manual drawing, or balls about a center reported previously. The proposed BML approach dissolves multiple testing through a multilevel model that more accurately accounts for data structure as well as shared information, and it consequentially improves inference efficiency. The modeling approach will be extended to other scenarios in our future work.

As a novel approach, BML here is applied to neuroimaging in dealing with multiplicity at the ROI level, with a potential extension to whole brain analysis in future work. We present this work in a purposefully (possibly overly) didactic style in the appendices, reflecting our own conceptual progression. Our goal is to convert the traditional voxel-wise GLM into an ROI-based BML through a step-wise progression of models (GLM ⟶ LME ⟶ BML). The paper is structured as follows. In the next section, we first formulate the population analysis at each ROI through univariate GLM (parallel to the typical voxel-wise population analysis), then turn multiple GLMs into one LME by pivoting the ROIs as the levels of a random-effects factor, and lastly convert the LME to a full BML. The BML framework does not make statistical inferences for each measuring entity (ROI in our context) in isolation. Instead, the BML weights and borrows the information based on the precision information across the full set of entities, striking a balance between data and prior knowledge; in a nutshell, the crucial feature here is that the ROIs, instead of being loose, are associated with each other through a Gaussianity assumption under BML. As a practical exemplar, we apply the modeling approach to an experimental dataset and compare its performance with the conventional univariate GLM. In the Discussion section, we elaborate the advantages, limitations, and prospects of BML in neuroimaging. Major acronyms and terms are listed in Table 1.

**Table 1.** Acronyms and terminology.

|  |  |  |  |
| --- | --- | --- | --- |
| BML | Bayesian multilevel | NHST | null hypothesis significance testing |
| FPR | false positive rate | NUTS | No-U-Turn sampler |
| FWE | family-wise error | power | chance of rejecting $H_0$ when $H_0$ is false |
| GLM | general linear model | PPC | posterior predictive check |
| HMC | Hamiltonian Monte Carlo | type I | chance of rejecting $H_0$ when $H_0$ is true (“false positive”) |
| LME | linear mixed-effects | type II | failing to reject $H_0$ when $H_0$ is false (“false negative”) |
| LOO | leave one out | type M | exaggerating the effect magnitude |
| MCMC | Monte Carlo Markov chain | type S | estimating the effect with an incorrect sign |

## Theory: Bayesian multilevel modeling

Throughout this article, the word *effect* refers to a quantity of interest, usually embodied in a regression (or correlation) coefficient, the contrast between two such quantities, or the linear combination of two or more such quantities from individual subject analysis. Italic letters in lower case (e.g., α) stand for scalars and random variables; lowercase, boldfaced italic letters (a) for column vectors; Roman and Greek letters for fixed and random effects in the conventional statistics context, respectively, on the righthand side of a model equation (the Greek letter θ is reserved for the effect of interest); *p(•)* represents a probability density function.

### Bayesian modeling for two-way random-effects ANOVA

As our main focus here is FMRI population analysis, we extend the BML approach for one-way ANOVA (Appendix D) to a two-way ANOVA structure, and elucidate the advantages of data calibration and partial pooling in more details. At the population level, the variability across n subjects has to be accounted for; in addition, the within-subject correlation structure among the ROIs also needs to be maintained. The conventional approach formulates r separate GLMs each of which fits the data *y_ij_* from the ith subject at the *j*th ROI,

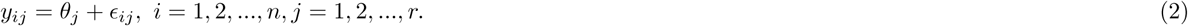

where *j = 1, 2,…, r, θ_j_* is the population effect at the *j*th ROI, and ∊_ij_ is the residual term that is assumed to independently and identically follow *N*(0, *σ^2^).* Each of the *r* models in (1) essentially corresponds to a Student’s t-test, and the immediate challenge is the multiple testing issue among those r models: with the assumption of exchangeability among the ROIs, is Bonferroni correction the only valid solution? If so, most neuroimaging studies would have difficulty in adopting ROI-based analysis due to this severe penalty, which may be the major reason that discourages the use of region-level analysis with a large number of regions. Alternatively, the *r* separate GLMs in (1) can be merged into one GLM by pooling the variances across the r ROIs,

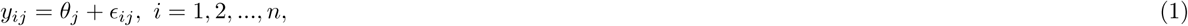

The two approaches, (1) and (2), usually render similar inferences unless the sampling variances are dramatically different across the ROIs. To compare different models through information criteria (Vehtari et al., 2017), we can solve the GLM (2) in a Bayesian fashion,

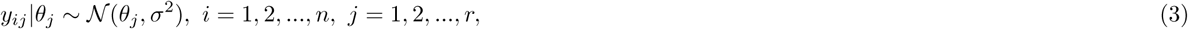

where the effects *θ_j_* are assigned with a noninformative prior so that no pooling is applied among the ROIs, leading to virtually identical inferences as the GLM (2).

We first extend one-way random-effects ANOVA (model (19) in Appendix D) to a two-way random-effects ANOVA, and formulate the following platform with data from *n* subjects,

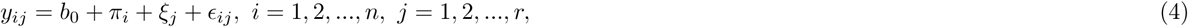

where *b_0_* represents the population effect, *π_i_* and *ξ_j_* code the deviation or random effect of the *i*th subject and *j*th ROI from the overall mean *b_0_*, respectively, and they are assumed to be *ii*d with *N*(0, λ^2^) and *N*(0, τ^2^), and ∊_ij_ is the residual term that is assumed to follow *N*(0, *σ^2^).*

Parallel to the situation with one-way ANOVA (Appendix D), the two-way ANOVA (4) can be conceptualized as an LME without changing its formulation. Specifically, the overall mean *b*_0_ is a fixed-effects parameter, while both the subject-and ROI-specific effects, *π_i_* and *ξ_j_*, are treated as random variables. In addition, we continue to define *θ_j_ = *b*_0_ + ξ_j_* as the effect of interest at the *j*th ROI. The LME framework has been well developed over the past half century, under which we can estimate variance components such as λ^2^ and τ^2^, and fixed effects such as *b*_0_ in (4). Therefore, conventional inferences can be made by constructing an appropriate statistic for a null hypothesis. Its modeling applicability and flexibility have been substantiated by its adoption in FMRI group analysis (Chen et al., 2013). Furthermore, the LME formulation (4) has a special layout, a crossed random-effects (or cross-classified) structure, which has been applied to inter-subject correlation (ISC) analysis for naturalistic scanning (Chen et al., 2017a) and to ICC analysis for ICC(2,1) (Chen et al., 2017c). A hierarchical model is a particular multilevel model in which parameters are nested within one another, and the cross-classified structure here showcases the difference between the two conceptions: the two clusters (ROI and subjects) intertwine with each other and form a factorial structure (*n* subjects by *r* ROIs), distinct from a hierarchical or nested one.

However, LME cannot offer a solution in making inferences regarding the ROI effects *θ_j_*: to estimate *θ_j_*, the LME (4) would become over-parameterized (i.e., an over-fitting problem). To proceed for the sake of intuitive interpretations, we temporarily assume a known sampling variance σ^2^, a known cross-subjects variance λ^2^, and a known cross-ROI variance τ^2^, and transform the ANOVA (4) to its Bayesian counterpart,

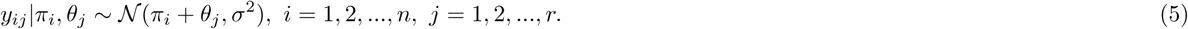

Then the posterior distribution of **θ_j_** with prior distributions, π_i_ ~ N(0, λ^2^) and **θ_j_** ~ *N(b_0_, τ^2^)*, can be analytically derived (Appendix E) with the data *y* = {*y_ij_*},

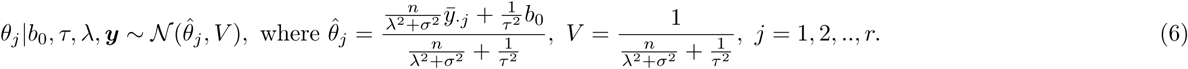

Similarly to the one-way ANOVA scenario (Appendix D), we have an intuitive interpretation for 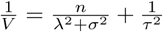: the posterior precision for *θ_j_*|b_0_, τ, λ, y is the sum of the cross-ROI precision 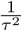 and the combined sampling precision 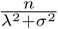. Under the r completely separate GLMs in (1), the cross-subjects variance λ^2^ and the sampling variance *σ^2^* could not be estimated separately. Interestingly, the following relationship,

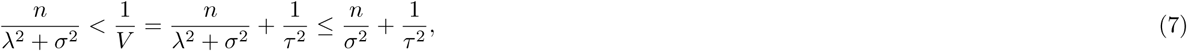

reveals that the posterior precision lies somewhere among the precisions of 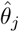 from the *r* separate GLMs. Furthermore, the posterior mode of 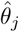 in (6) can be expressed as a weighted average between the individual sample means 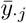 and the overall mean *b_0_*,

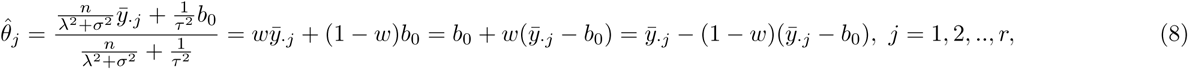

where the weight 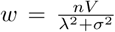, indicating the counterbalance of partial pooling between the individual mean 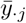 for the *j*th entity and the overall mean *b*_0_, the adjustment of **θ_j_** from the overall mean *b*_0_ toward the observed mean 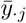, or the observed mean 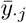 being shrunk toward the overall mean *b*_0_.

Related to the concept of ICC, the correlation between two ROIs, *j_i_* and *j*_2_, due to the fact that they are measured from the same set of subjects, can be derived in a Bayesian fashion as,

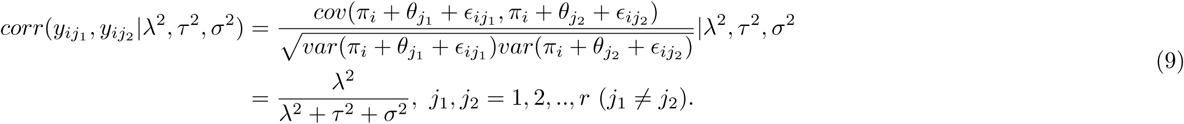

Similarly, the correlation between two subjects, *i_1_* and *i_2_*, due to the fact that their effects are measured from the same set of ROIs, can be derived in a Bayesian fashion as,

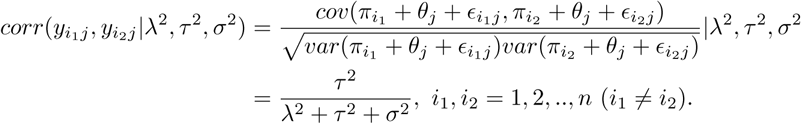

The exchangeability assumption is crucial here as well for the BML system (4). Conditional on ξ_j_ (i.e., when the ROI is fixed at index *j*), the subject effects *π_i_* can be reasonably assumed to be exchangeable since the experiment participants are usually recruited randomly from a hypothetical population pool as representatives (thus the concept of coding them as dummy variables). As for the ROI effects ξ_j_, here we simply assume the validity of exchangeability conditional on the subject effect *nπ_i_* (i.e., when subject is fixed at index *i*), and address the validity later in Discussion.

To summarize, the main difference between the conventional GLM and BML lies in the assumption about the brain regions: the effects (e.g., **θ_j_** in (3)) are assumed to have a noninformative flat prior while they are assigned with a Gaussian prior under BML. In other words, the effect at each region is estimated independently from other regions under GLM, thus there is no information shared across regions. In contrast, the effects across regions are shared, regularized and partially pooled through the Gaussian assumption under BML for the effects across regions; such a cross-region Gaussianity bears the same rationale as the cross-subject Gaussianity. So far, we have presented a “simplest” BML scenario. Specifically, we have: ignored the possibility of incorporating any explanatory variables such as subject-specific quantities (e.g., age, IQ) or behavioral data (e.g., reaction time); assumed known variances such as τ^2^ and σ^2^; and presumed that the data *y_ij_* have been directly measured without precision information available. Further extensions are needed and discussed for realistic applications in the next subsection.

### Further extensions of Bayesian modeling for two-way random-effects ANOVA and full Bayesian implementations

To gain intuitive interpretations, we have so far assumed that the variances σ^2^, λ^2^ and τ^2^ in (5) (and *σ^2^* in (23) of Appendix D) are known. In practice, those parameters for the prior distributions are not available. Approximate (or empirical) Bayesian approaches could be adopted to provide a computationally economical “workaround” solution. For example, one possibility is to first solve the corresponding LME and directly apply the estimated variances to the analytical formula (6) (and (24) in Appendix D). However, there are two limitations associated with approximate Bayesian approaches. The reliability or uncertainty for the estimated variances are not taken into consideration and thus may result in inaccurate posterior distributions. In addition, analytical formulas such as (6) (and (24) in Appendix D) are usually not available when we extend the prototypical models (5) (and (23) in Appendix D) to more generic scenarios, as shown below.

At the population level, one may incorporate one or more subject-specific covariates such as subject-grouping variables (patients vs. controls, genotypes, adolescents vs. adults) and/or quantitative explanatory variables (age, behavioral or biometric data). To be able to adapt such scenarios, we first need to expand the models considered previously with a simple intercept (Student’s t-test) to r separate GLMs and one GLM with pooled variances, generalizing the models (1) and (3), respectively, to

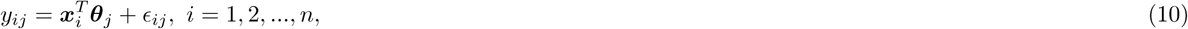

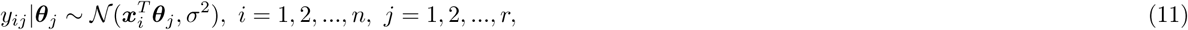

where the vector *x_i_* contains the subject-specific values of the covariates, with its first component 1 that is associated with the intercept, and the vector **θ_j_** codes the effects associated with the covariates x_i_ (and each component in **θ_j_** is assigned with a noninformative prior in (11)), *j* = 1, 2, …, *r*. In parallel, the conventional two-way random-effects ANOVA or LME (4) evolves to

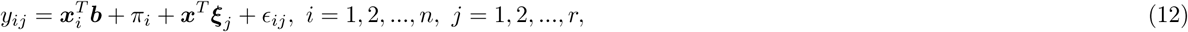

where b and ξ_j_ represent the population effects and subject-wise deviations corresponding to those covariates, respectively. Similarly, the BML counterpart can be formulated as

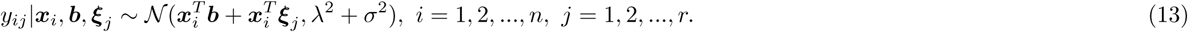

Under the BML (13), the effect of interest **θ_j_** can be an element of *b*, the intercept (as in (5)) or the effect for one of the covariates x_i_. Similar to models with varying intercepts such as (4) and (5), both intercepts and slopes are assumed to be different across ROIs in the models (12) and (13), and they are usually referred to as models with varying intercepts and slopes. When there is only one covariate *x_i_,* the four models (10), (11), (12) and (13) simplify to, respectively,

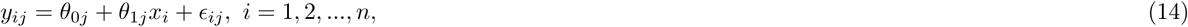

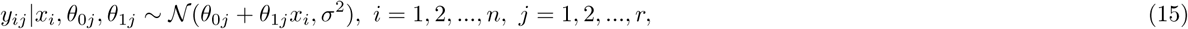

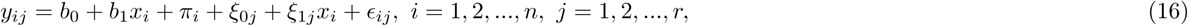

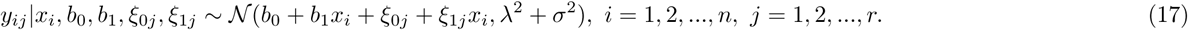

The discussion so far has assumed that data *y_ij_* are directly collected without measurement errors. However, in some circumstances (including neuroimaging) the data are summarized through one or more analytical steps. For example, the data y_ij_ in FMRI can be the BOLD responses from subjects under a condition or task that are estimated through a time series regression model, and the estimates are not necessarily equally reliable. Therefore, a third extension is desirable to broaden our model (13) so that we can accommodate the situation where the separate variances 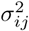 of measurement errors for each ROI and subject are known and should be included in the model (13) as inputs, instead of being treated as one hyperparameter. Similarly to the conventional meta-analysis, a BML with known sampling variances can be effectively analyzed by simply treating the variances as known values.

### Numerical implementations of BML

Since no analytical formula is generally available for the BML (13), we proceed with the full Bayesian approach hereafter, and adopt the algorithms implemented in Stan, a probabilistic programming language and a math library in C++ on which the language depends (Stan Development Team, 2017). In Stan, the main engine for Bayesian inferences is No-U-Turn sampler (NUTS), a variant of Hamiltonian Monte Carlo (HMC) under the category of gradient-based Markov chain Monte Carlo (MCMC) algorithms.

Some conceptual and terminological clarifications are warranted here. Under the LME framework, the differentiation between fixed- and random-effects is clearcut: fixed-effects parameters (e.g., b in (12)) are considered universal constants at the population level to be estimated; in contrast, random-effects variables (e.g., *π_i_* and ξ_j_ in (12)) are assumed to be varying and follow a presumed distribution. However, there is no such distinction between fixed and random effects in Bayesian formulations, and all effects are treated as parameters and are assumed to have prior distributions. Nevertheless, there is a loose correspondence between LME and BML: fixed effects under LME are usually termed as population effects under BML, while random effects in LME are typically referred to as entity effects^1^ under BML.

Essentially, the full Bayesian approach for the BML systems (5) and (13) can be conceptualized as assigning hyperpriors to the parameters in the LME or ANOVA counterparts (4), and (12). Our prior distribution choices follow the general recommendations in Stan (Stan Development Team, 2017). Regarding hyperpriors, an improper flat (noninformative uniform) distribution over the real domain for the population parameters (e.g., b in (13)) is adopted, since we usually can afford the vagueness thanks to the usually satisfactory amount of information available in the data at the population level. For the scaling parameters at the entity level, the variances for the cross-subjects effects π_i_ and as well as in the variance-covariance matrix for ξ_j_ in (13), we use a weakly informative prior such as a Student’s half-t(3,0,1)^2^ or half-Gaussian *N*(0,1) (restricting to the positive half of the respective distribution). For the covariance structure of, the LKJ correlation prior^3^ is used with the parameter ζ = 1 (i.e., jointly uniform over all correlation matrices of the respective dimension). Lastly, the variance for the residuals ∊_ij_ is assigned with a half Cauchy prior with a scale parameter depending on the standard deviation of y_ij_. To summarize, besides the Bayesian framework under which hyperpirors provide a computational convenience through numerical regularization, the major difference between BML and its univariate GLM counterpart is the Gaussian assumption for the ROIs (e.g., *θ_j_* ~ N(b_0_,τ^2^) in the model (5)) that plays the pivotal role of pooling and sharing the information among the brain regions. It is this partial pooling that effectively takes advantage of the effect similarities among the ROIs and achieves higher modeling efficiency.

Another different aspect about Bayesian inference is that it hinges around the whole posterior distribution of an effect. For practical considerations in results reporting, modes such as mean and median are typically used to show the centrality, while a quantile-based (e.g., 95%) interval or highest posterior density provides a condensed and practically useful summary of the posterior distribution. The typical workflow to obtain the posterior distribution for an effect of interest is the following. Multiple (e.g., 4) Markov chains are usually run in parallel with each of them going through a predetermined number (e.g., 2000) of iterations, half of which are thrown away as warm-up (or “burn-in”) iterations while the rest are used as random draws from which posterior distributions are derived. To gauge the consistency of an ensemble of Markov chains, the split 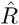 statistic (Gelman et al., 2014) is provided as a potential scale reduction factor on split chains and as a diagnostic parameter to assist the analyst in assessing the quality of the chains. Ideally, fully converged chains correspond to 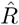 = 1.0, but in practice 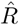 < 1.1 is considered acceptable. Another useful parameter, the number of effective sampling draws after warm-up, measures the number of independent draws from the posterior distribution that would be expected to produce the same standard deviation of the posterior distribution as is calculated from the dependent draws from HMC. As the sampling draws are not always independent with each other, especially when Markov chains proceed slowly, one should make sure that the effective sample size is large enough relative to the total sampling draws so that a reasonable accuracy can be achieved to derive the quantile intervals for the posterior distribution. For example, a 95% quantile interval requires at least an effective sample size of 100. As computing parallelization can only be executed for multiple chains of the HMC algorithms, the typical BML analysis can be effectively conducted on any system with at least 4 CPUs.

One important aspect of the Bayesian framework is model quality check through various prediction accuracy metrics. The aim of the quality check is not to reject the model, but rather to check whether it fits the data well. For instance, posterior predictive check (PPC) simulates replicated data under the fitted model and then graphically compares actual data *y_ij_* to the model prediction. The underlying rationale is that, through drawing from the posterior predictive distribution, a reasonable model should generate new data that look similar to the acquired data at hand. As a model validation tool, PPC intuitively provides a visual tool to examine any systematic differences and potential misfit of the model, similar to the visual examination of plotting a fitted regression model against the original data. Leave-one-out (LOO) cross-validation using Pareto-smoothed importance sampling (PSIS) is another accuracy tool (Vehtari et al., 2017) that uses probability integral transformation (PIT) checks through a quantile-quantile (Q-Q) plot to compare the LOO-PITs to the standard uniform or Gaussian distribution.

### BML applied to an ROI-based group analysis

To demonstrate the performances of BML in comparison to the conventional univariate approach at the ROI level, we utilized an experimental dataset from a previous FMRI study (Xiao et al., 2018). Briefly, a cohort of 124 typically developing children (mean age = 6.6 years, SD = 1.4 years, range = 4 to 8.9 years; 54 males) was scanned while they watched Inscapes, a movie paradigm designed for collecting resting-state data to reduce potential head motion. In addition, a subject-level covariate was included in the analysis: the overall theory of mind ability based on a parent-report measure (the theory of mind inventory, or ToMI). FMRI images were acquired with the following EPI scan parameters: *b*_0_ = 3 T, flip angle = 70 °, echo time = 25 ms, repetition time = 2000 ms, 36 slices, planar field of view = 192 x 192 mm^2^, voxel size = 3.0 x 3.0 x 3.5 mm^3^, 210 volumes with a total scanning time of 426 seconds. Twenty-one ROIs (Table 3) were selected from the literature because of their potential relevancy to the current study, and they were neither chosen nor defined per the whole brain analysis results of the current data. Mean Fisher-transformed z-scores were extracted at each ROI from the output of seed-based correlation analysis (seed: right temporo-parietal junction at the MNI coordinates of (50, −60, 18)) from each of the 124 subjects. The effect of interest at the population level is the relationship at each brain region between the behavioral measure of the overall ToMI and the region’s association with the seed. A whole brain analysis showed the difficulty of some clusters surviving FWE correction (Table 2).

**Table 2.** ROIs and FWE correction for their associated clusters^a^

| voxel-wise $p$ | cluster threshold | number of surviving ROIs | ROIs |
| --- | --- | --- | --- |
| 0.001 | 28 | 2 | R PCC, PCC/PrC |
| 0.005 | 66 | 4 | R PCC, PCC/PrC., L IPL, L TPJ |
| 0.01 | 106 | 4 | R PCC, PCC/PrC., L IPL, L TPJ |
| 0.05 | 467 | 4 | R PCC, PCC/PrC., L IPL, L TPJ |
| 0.05* | 467 | (4) | (L aMTS/aMTG, R TPJp, vmPFC, dmPFC) |

<sup>a</sup>Monte Carlo simulations were conducted using a mixed exponential spatial autocorrelation function (Cox et al., 2017) instead of FWHM to determine the cluster threshold (voxel size: $3 \times 3 \times 3 \text{ mm}^3$ ). The ROI abbreviations are listed in Table 3.
\*Special note for the last row (voxel-wise $p$ -value of 0.05): four ROIs including L IPL, L TPJ, R PCC, PCC/PrC survived together with their clusters from the FWE correction, and the other four ROIs listed here (L aMTS/aMTG, R TPJp, vmPFC, and dmPFC) did not survive with their clusters but showed some evidence of effect when the cluster size requirement was dropped.

**Table 3.** MNI coordinates of the 21 ROIs^a^

| No | ROI | Coordinates ($x, y, z$) |
| --- | --- | --- |
| 1 | R PCC | (8, -59, 35) |
| 2 | R TPJp | (56, -56, 25) |
| 3 | R Insula | (49, -8, -11) |
| 4 | L IPL | (-55, -65, 27) |
| 5 | L SFG | (-7, 58, 21) |
| 6 | R IFG (BA45) | (47, 22, 6) |
| 7 | R IFG (BA9) | (60, 25, 19) |
| 8 | L MTG | (-51, -62, 5) |
| 9 | L CG | (-5, 8, 42) |
| 10 | L IFG | (-46, 24, 7) |
| 11 | ACC | (0, 38, 10) |
| 12 | SGC | (-2, 32, -8) |
| 13 | PCC/PrC | (-2, -52, 26) |
| 14 | dmPFC | (-2, 5, 14) |
| 15 | L TPJ | (-46, -66, 18) |
| 16 | L vBG | (-6, 10, -8) |
| 17 | R vBG | (6, 10, -8) |
| 18 | L aMTS/aMTG | (-54, -10, -20) |
| 19 | R Amy/Hippo | (24, -8, -22) |
| 20 | L Amy/Hippo | (-24, -10, -20) |
| 21 | vmPFC | (-2, 50, -10) |

<sup>a</sup>The 21 ROIs were chosen because of their potential involvement for the current experiment based on previous studies. Each ROI was created as a ball with a center at the coordinates (in millimeters) from the literature (Xiao et al., 2017) and a radius of 6 mm. ROI abbreviations: L, left hemisphere; R, right hemisphere; PCC/PrC, precuneus/posterior cingulate cortex; TPJp, posterior temporo-parietal junction; IPL, inferior parietal lobe; SFG, superior frontal gyrus; IFG, inferior frontal gyrus; aMTS/aMTG, anterior middle temporal sulcus/gyrus; CG, cingulate gyrus; ACC, anterior cingulate cortex; SGC, subgenual cingulate cortex; dmPFC, dorsomedial prefrontal cortex; vBG, ventral basal ganglia; Amy/Hippo, amygdala/hippocampus; vmPFC, ventromedial prefrontal cortex.

The data from the 21 ROIs were analyzed through the modeling triplets, GLMs (14) and (15), LME (16) and BML (17), with the effect of interest at the *j*th ROI being the relationship between ToMI and the ROI’s association with the seed: *θ_1j_* = *b_1_ + ξ_1j_*. The exchangeability assumption for LME and BML was deemed reasonable because, prior to the analysis, no specific information was available regarding the order and relatedness of the effects across subjects and ROIs. It is worth noting that the data were skewed with a longer right tail than left (black solid curve in Fig. 3a and Fig. 3b).

**Figure 1:**
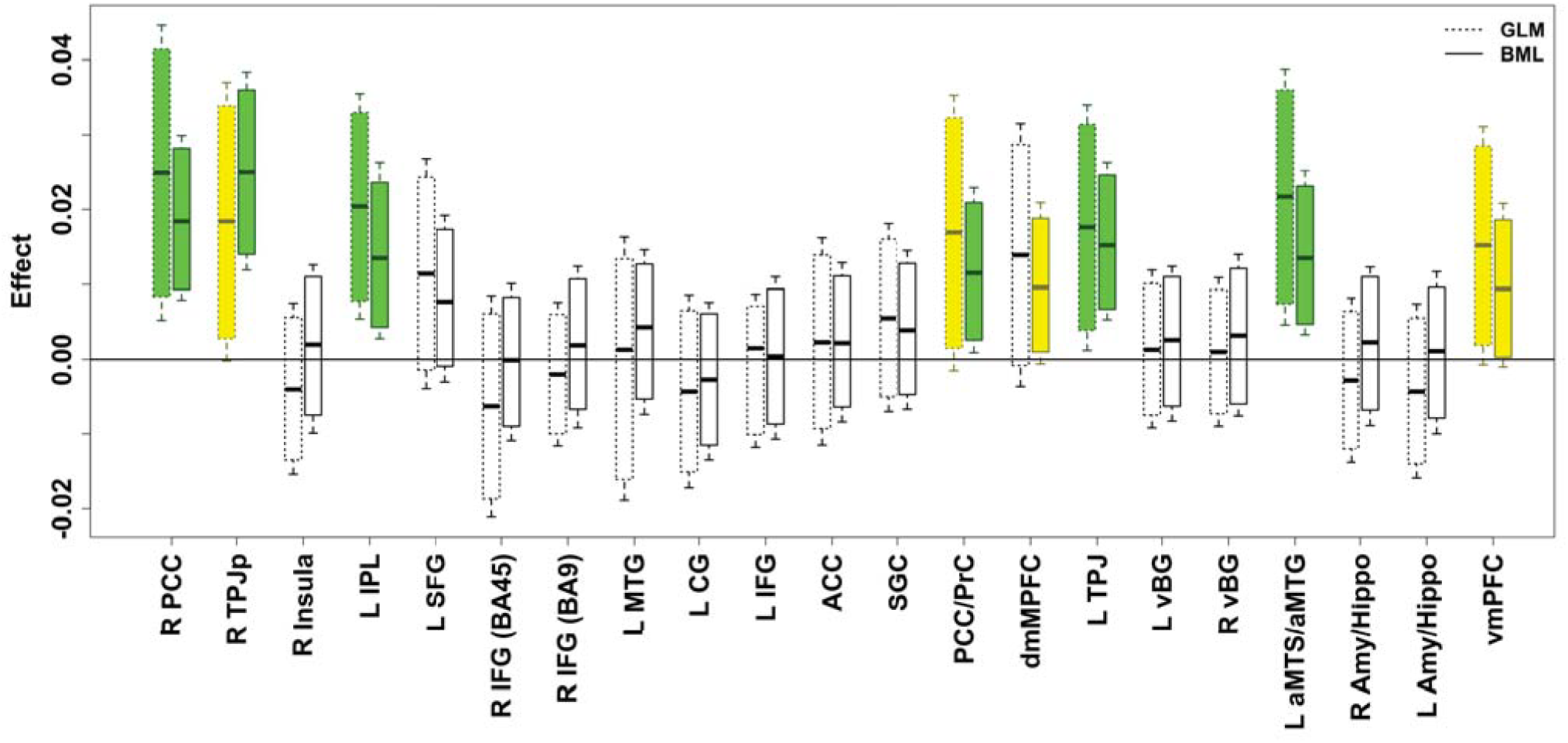
Comparisons of results between the conventional GLM and BML

**Figure 2:**
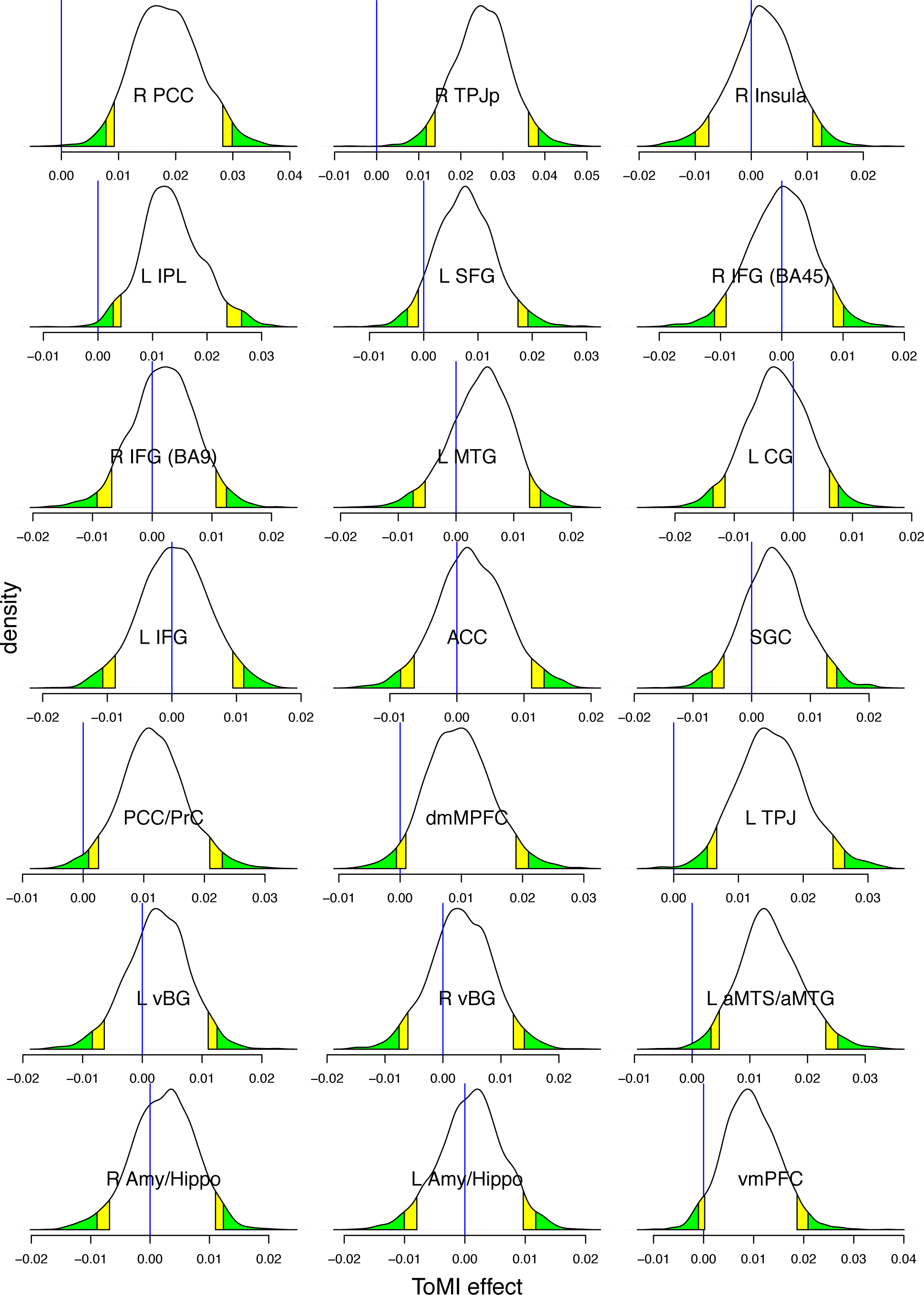
Posterior density distributions based on 2000 draws from BML. The vertical blue line indicates zero ToMI effect, yellow and green tails mark the 90% and 95% quantile intervals, respectively, and the ROIs with strong evidence of ToMI effect can be identified as the blue line being within the color tails. Compared to the conventional confidence interval that is flat and inconvenient to interpret, the posterior density provides much richer information about each effect such as spread, shape and skewness.

**Figure 3:**
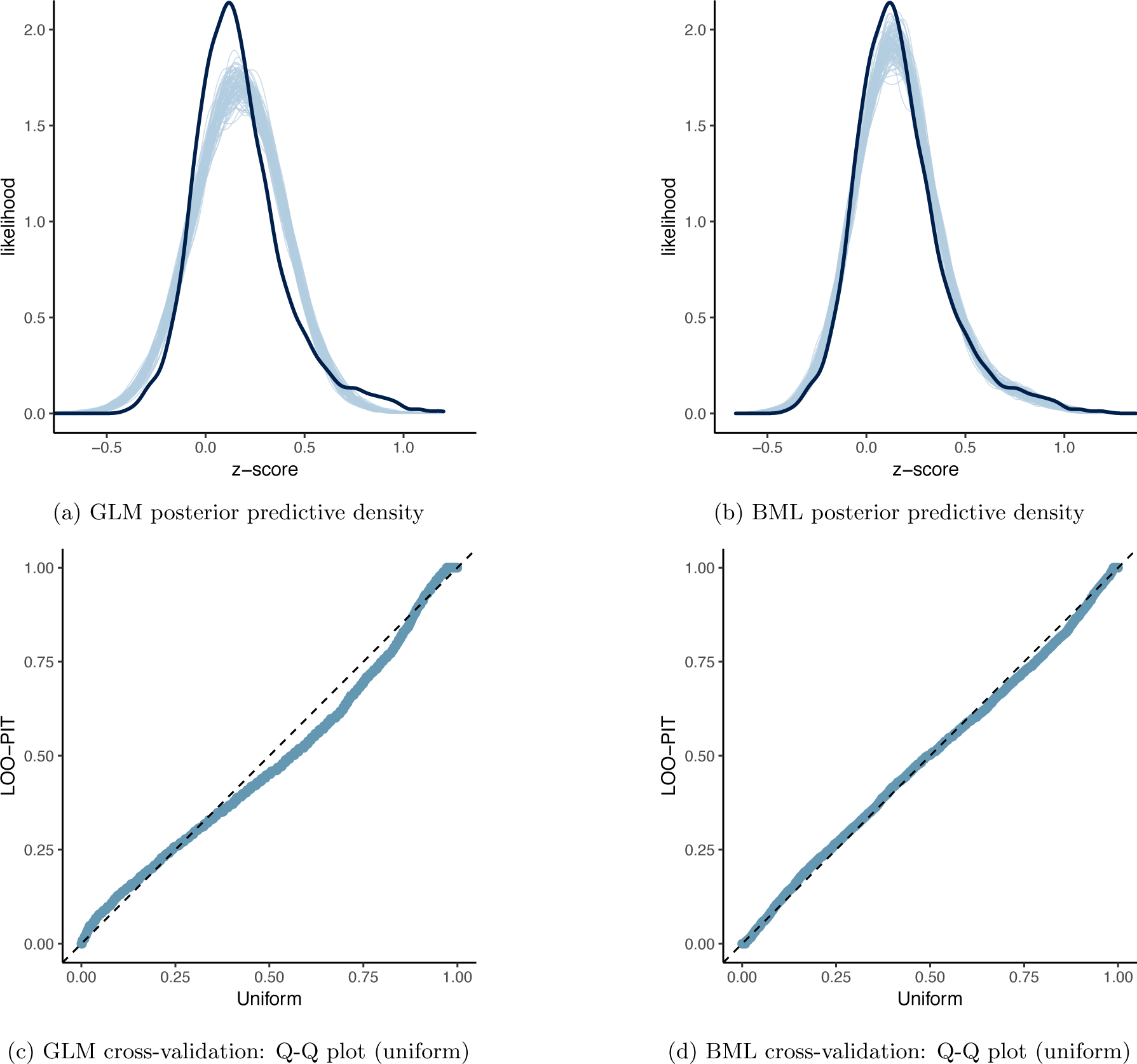
Model performance comparisons through posterior predictive check between conventional univariate GLM (a and c) and BML (b and d). The subfigures *a* and *b* show the posterior predictive density overlaid with the raw data from the 124 subjects at the 21 ROIs for GLM and BML, respectively: solid black curve is the raw data at the 21 ROIs with linear interpolation while the fat curve in light blue is composed of 100 sub-curves each of which corresponds to one draw from the posterior distribution based on the respective model. The differences between the two curves indicate how well the respective model fits the raw data. BML fitted the data better than GLM at the peak and both tails as well as the skewness because pooling the data from both ends toward the center through shrinkage clearly validates our adoption of BML. The subfigures c and d contrast GLM and BML through cross-validation with leave-one-out log predictive densities through the calibration of marginal predictions from 100 draws; the calibration is assessed by comparing probability integral transformation (PIT) checks to the standard uniform distribution. The diagonal dished line indicates a perfect calibration: there are some suboptimal calibration for both models, but BML is clearly a substantial improvement over GLM. To simulate the posterior predictive data for the conventional ROI-based approach *(a* and c), the Bayesianized version of GLM (15) was adopted with a noninformative uniform prior for the population parameters.

When fitted at each ROI separately with GLM (simple regression in this case) using the overall ToMI as an explanatory variable, the model yielded lackluster fitting (Fig. 3a) in terms of skewness, the two tails, and the peak area. As shown in Fig. 1, five ROIs (R PCC, R TPJp, L IPL, L TPJ, and L aMTS/aMTG) reached a two-tailed significance level of 0.05, and two ROIs (PCC/PrC and vmPFC) achieved a two-tailed significance level of 0.1 (or one-tailed significance level of 0.05 if directionality was *a priori* known). However, the burden of FWE correction (e.g., Bonferroni) for the ROI-based approach with univariate GLM is so severe that none of the ROIs could survive the penalizing metric.

The ROI data were fitted with LME (16) and BML (17) using the overall ToMI as an explanatory variable through, respectively, the R (R Core Team, 2017) package lme4 (Bates et al., 2015) and Stan with the code translated to C++ and compiled. Runtime for BML was 5 minutes including approximately 1 minute of code compilation on a Linux system (Fedora 25) with AMD Opteron 6376 at 1.4 GHz. All the parameter estimates at the population level were quite similar between the two models (Table 4), indicating that the typical hyperpriors adopted for BML had little impact on parameter estimation. However, of interest here are the effects at the entity (i.e., ROI), not population, level, which could be derived through BML but not LME. As for those effects at the ROI level, compared to the traditional ROI-based GLM, the shrinkage under BML can be seen in Fig. 1: most effect estimates were dragged toward the center. Similar to the ROI-based GLM without correction, BML demonstrated (Figures 1 and 2) strong evidence within 95% quantile interval of the overall ToMI effect at six ROIs (R PCC, R TPJp, L IPL, PCC/PrC, L TPJ, and L aMTS/aMTG), and within 90% (or 95% if directionality was *a priori* known) quantile interval at two additional ROIs (dmMPFC and vmPFC).

**Table 4.**
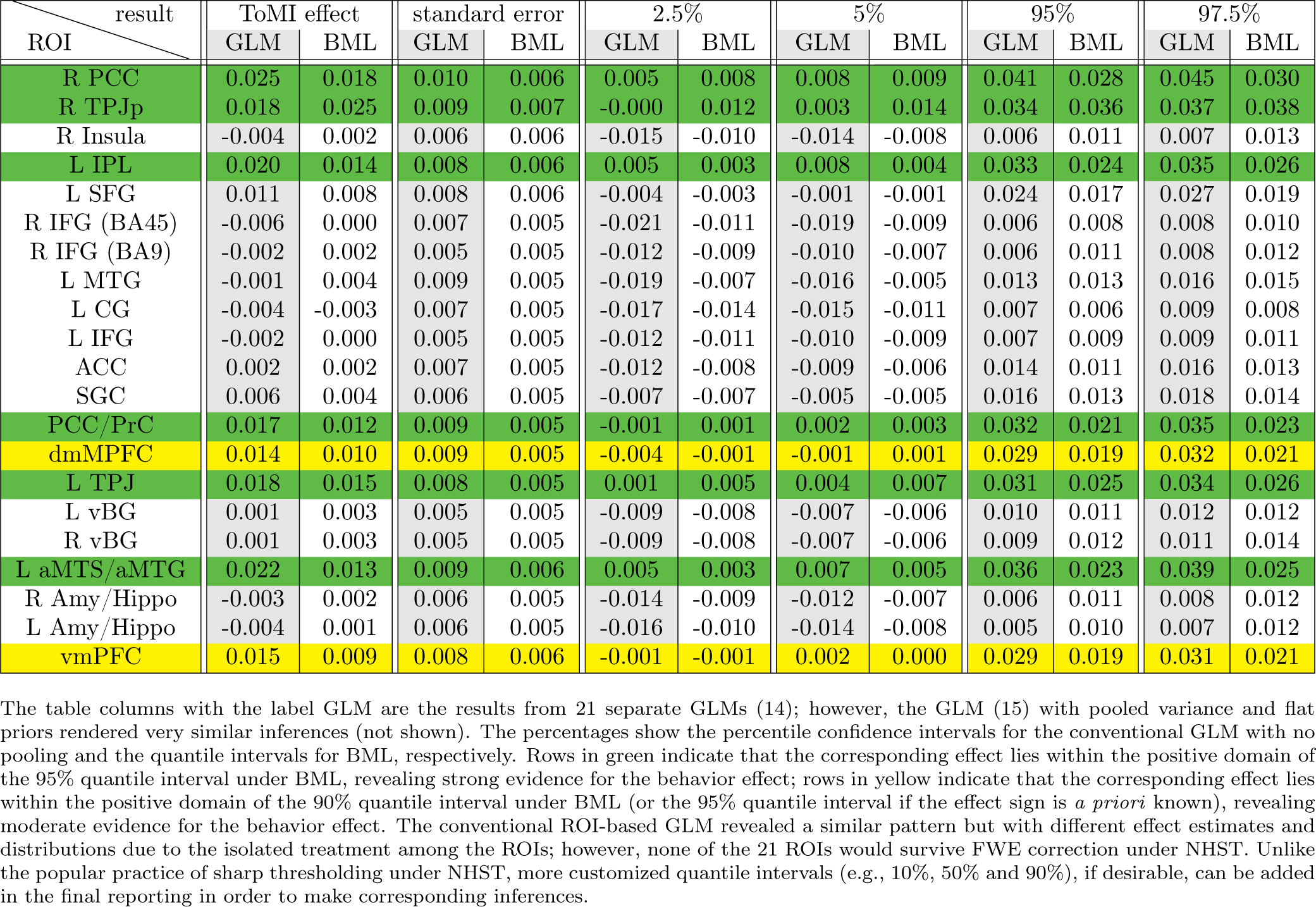
Comparisons among GLM, LME and BML. The seed-based correlation results at 21 ROIs from 124 subjects were fitted with LME (using R package lme4) and BML (using Stan with 4 chains and 1,000 iterations) separately, in which overall ToMI was an explanatory variable. Random effects under LME correspond to group/entity-level effects plus family specific parameters (variance σ^2^ for residuals *∊_ij_*) under BML, while fixed effects under LME correspond to population-level effects under BML. The parameter estimates from the LME output (a) and the BML output (b) are very similar, even though priors were injected into BML. All *R* values under BML were less than 1.1, indicating that all the 4 chains converged well. The effective sizes for the population- and group/entity-level effect of ToMI were 468 and 947, respectively, enough to warrant quantile accuracy in summarizing the posterior distributions. To directly compare with BML, the Bayesianized version of GLM (15) was fitted with the data, and the higher predictive accuracy of BML is seen in (c) with its substantial lower out-of-sample deviance measured by the leave-one-out information criterion (LOOIC), the widely applicable (or Watanabe-Akaike) information criterion (WAIC) through leave-one-out cross-validation, and the corresponding standard error (SE).

One exception to the general shrinkage under BML is that the median effect, 0.025, at the region of R TPJp (second row in the table and box plot of Fig. 1) was actually higher than that under GLM, 0.018. Such an exception occurred because the final result is a combination or a tug of war between the shrinkage impact as shown in (8) and the correlation structure among the ROIs as shown in (9). Noticeably, the quality and fitness of BML can be diagnosed and verified through posterior predictor check (Fig. 3a and Fig. 3b) that compares the observed data with the simulated data based on the model: not only did BML accommodate the skewness of the data better than GLM, but also did the partial pooling render much better fit for the peak and both tails as well. Cross validation through LOO (Table 4c, Fig. 3c and Fig. 3d) also manifested the advantage of BML fitting over GLM. Nevertheless, there is still room for the improvement of BML: the peak area could be fitted better, which may require nonlinearity or incorporating other potential covariates.

One apparent aspect that the ROI-based BML excels is the completeness and transparency in results reporting: if the number of ROIs is not overwhelming (e.g., less than 100), the summarized results for every ROI can be completely presented in a tabular form *(c.f.* Fig. 1) and in full distributions of posterior density (Fig. 2). It is worth emphasizing that Bayesian inferences focus less on the point estimate of an effect and its associated quantile interval, but more on the whole posterior density as shown in Fig. 2 that offers more detailed information about the effect uncertainty. Unlike the whole brain analysis in which the results are typically reported as the tips of icebergs above the water, posterior density reveals the spread, shape and skewness regardless of the statistical evidence. In addition, one does not have to stick to a single harsh thresholding when deciding a criterion on the ROIs for discussion; for instance, even if an ROI lies outside of, but close to, the 95% quantile interval (e.g., dmMPFC and vmPFC in Figures 1 and 2), it can still be reported and discussed as long as all the details are revealed. Such flexibility and transparency are difficult to navigate or maneuver through cluster thresholding at the whole brain level. As a counterpart to NHST, a probability metric could still be provided for each effect under BML in the sense as illustrated in Table 5; however, we opt not to do so for two reasons: 1) such a probability measure could be easily misinterpreted in the NHST sense, and, more importantly, 2) it is the predictive intervals shown in Fig. 1 and the complete posterior distributions illustrated in Fig. 2, not the single probabilities, that fully characterize the posterior distribution, providing richer information than just binary (“in or out”) thresholding.

**Table 5.**
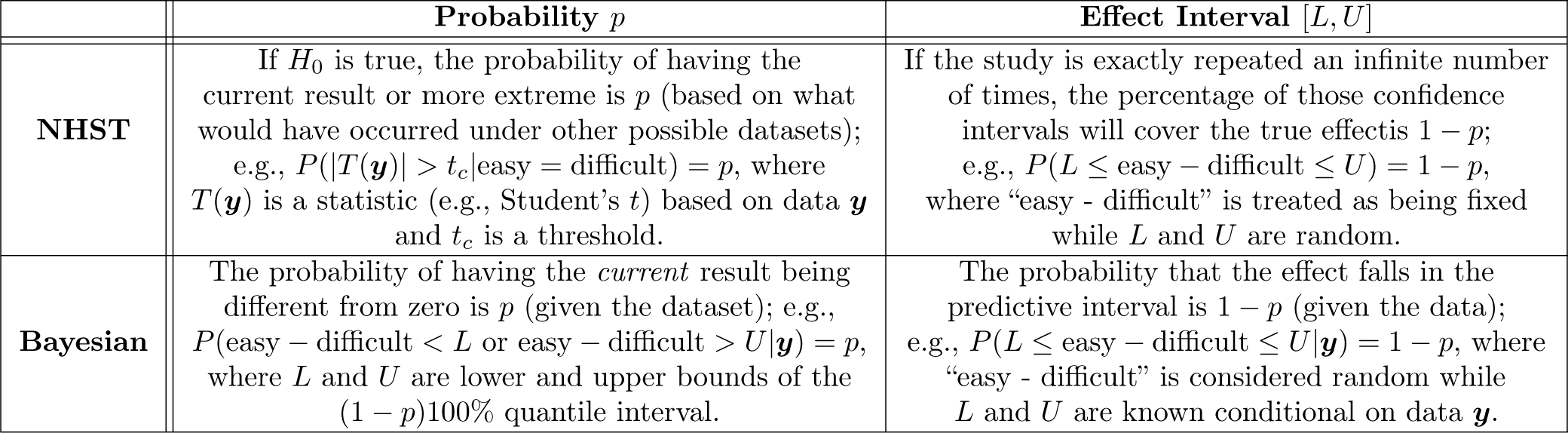
Interpretation differences between NHST and Bayesian framework

Interestingly, those four regions (L IPL, L TPJ, R PCC, PCC/PrC) that passed the FWE correction at the voxel-wise p-cutoff of 0.005 (Table 2) in the whole brain analysis were confirmed with the ROI-based BML (Figures 1 and 2). Moreover, another four regions (L aMTS/aMTG, R TPJp, vmPFC, dmPFC) revealed some evidence of ToMI effect under BML. In contrast, these four regions did not stand out in the whole brain analysis after the application of FWE correction at the cluster level regardless of the voxel-wise p-threshold (Table 2), even though they would have been evident if the cluster size requirement were not as strictly imposed.

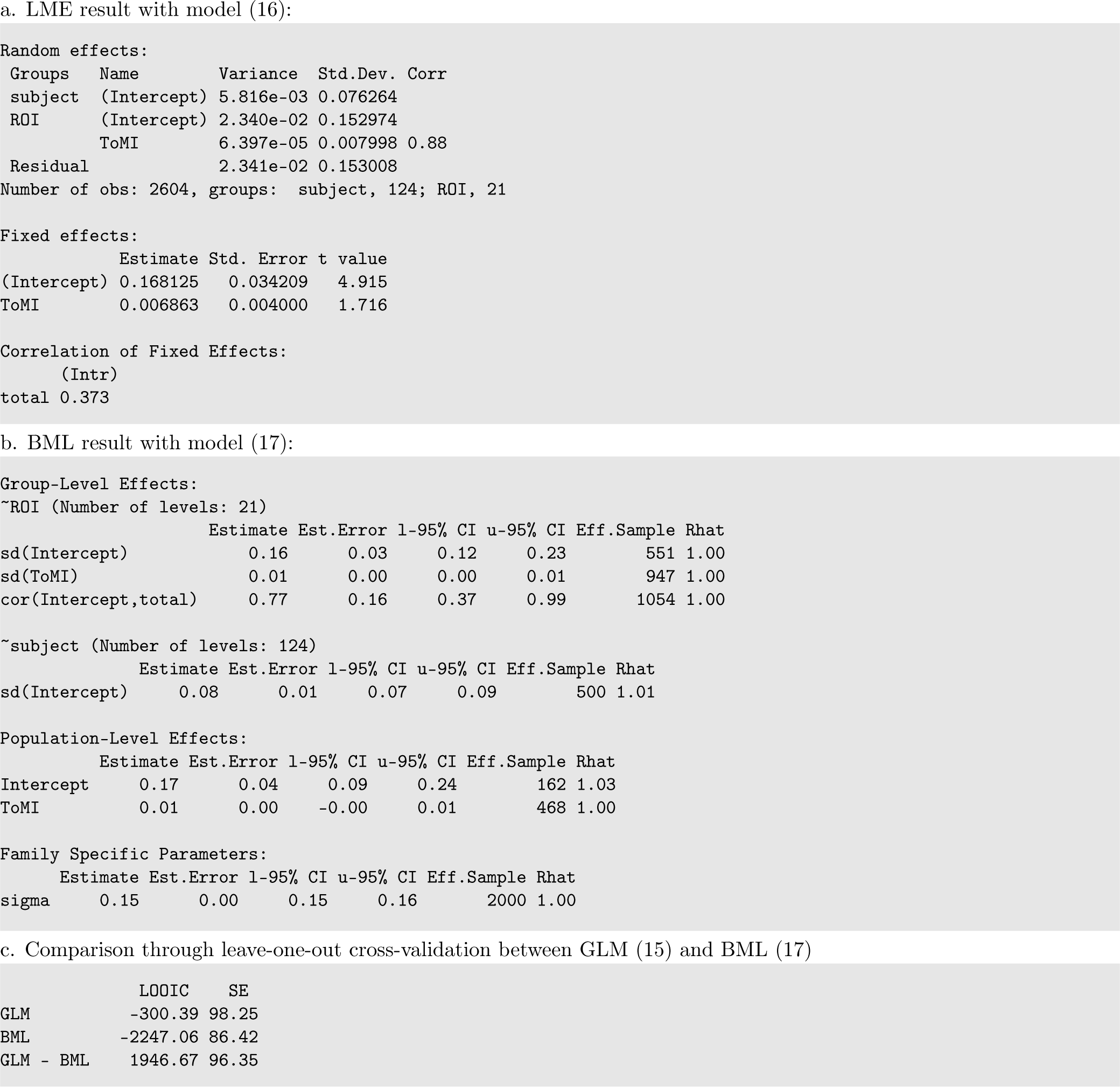

## Discussion

### Current approaches to correcting for FPR

Arbitrariness is involved in the multiple testing correction of parametric methods. In the conventional statistics framework, the thresholding bar ideally plays the role of winnowing the wheat (true effect^4^) from the chaff (random noise), and a *p*-value of 0.05 is commonly adopted as a benchmark for comfort in most fields. However, one big problem facing the correction methods for multiple testing is the arbitrariness surrounding the thresholding, in addition to the arbitrariness of 0.05 itself. Both Monte Carlo simulations and random field theory start with a voxel-wise probability threshold (e.g., 0.01,0.005, 0.001) at the voxel (or node) level, and a spatial threshold is determined in cluster size so that overall FPR can be properly controlled at the cluster level. If clusters are analogized as islands, each of them may be visible at a different sea level (voxel-wise *p*-value). As the cluster size based on statistical filtering plays a leveraging role, with a higher statistical threshold leading to a smaller cluster cutoff, a neurologically or anatomically small region can only gain ground with a low p-value while large regions with a relatively large p-value may fail to survive the criterion. Similarly, a lower statistical threshold (higher *p*) requires a higher cluster volume, so smaller regions have little chance of reaching the survival level. In addition, this arbitrariness in statistical threshold at the voxel level poses another challenge for the investigator: one may lose spatial specificity with a low statistical threshold since small regions that are contiguous may get swamped by the overlapping large spatial extent; on the other hand, sensitivity may have to be compromised for large regions with low statistic values when a high statistical threshold is chosen. A recent critique on the rigor of cluster formation through parametric modeling (Eklund et al., 2016) has resulted in a trend to require a higher statistical thresholding bar (e.g., with the voxel-wise threshold below 0.005 or even 0.001); however, the arbitrariness persists because this trend only shifts the probability threshold range.

Permutation testing has its share of arbitrariness in multiple testing correction too. As an alternative to parametric methods, an early version of permutation testing (Nichols and Holmes, 2001) bears similar arbitrary issues. It starts with the construction of a null distribution through permutations in regard to a maximum statistic (either maximum testing statistic or maximum cluster size based on a predetermined threshold for the testing statistic). The original data are assessed against the null distribution, and the top winners at a designated rate (e.g., 5%) among the testing statistic values or clusters will be declared as the surviving ones. While the approach is effective in maintaining the nominal FWE level, two problems are embedded with the strategy. First of all, the spatial properties are not directly taken into consideration in the case of maximum testing statistic. For example, an extreme case to demonstrate the spatial extent issue is that a small cluster (or even a single voxel) might survive the permutation testing as long as its statistic value is high enough (e.g., t(20) = 6.0) while a large cluster with a relatively small maximum statistic value (e.g., t(20) = 2.5) would fail to pass the filtering. The second issue is the arbitrariness involved in the primary thresholding for the case of maximum cluster size: a different primary threshold may end up with a different set of clusters. That is, the reported results may likely depend strongly on an arbitrary threshold.Addressing these two problems, a later version of permutation testing (Smith and Nichols, 2009) takes an integrative consideration between signal strength and spatial relatedness, and thus solves both problems involving the earlier version of permutation testing^5^. Such an approach has been implemented in programs such as Randomise and PALM in FSL using threshold-free cluster enhancement (TFCE) (Smith and Nichols, 2009) and in 3dttest++ in AFNI using equitable thresholding and clusterization (ETAC) (Cox, 2018). Nevertheless, the adoption of permutations in neuroimaging, regardless of the specific version, is not directly about the concern of distribution violation as in the classical nonparametric setting (in fact, a pseudo-t value is still computed at each voxel in the process); rather, it is the randomization among subjects in permutations that creates a null distribution against which the original data can be tested at the whole brain level.

We argue that spatial size as a correction leverage unnecessarily pays the cost of lower identification power to achieve the nominal false positive level. All of the current correction methods, parametric and nonparametric, are still meant to use spatial extent or the combination of spatial extent and signal strength as a filter to control the overall FPR at the whole brain level. They all share the same hallmark of sharp thresholding at a preset acceptance level (e.g., 5%) under NHST, and they all use spatial extent as a leverage, penalizing regions that are anatomically small and rewarding large smooth regions (Cremers et al., 2017). Due to the unforgiving penalty of correction for multiple testing, some workaround solutions have been adopted by focusing the correction on a reduced domain instead of the whole brain. For example, the investigator may limit the correction domain to gray matter or regions based on previous studies. Putting the justification for these practices aside, accuracy is a challenge in defining such masks; in addition, spatial specificity remains a problem, shared by the typical whole brain analysis, although to a lesser extent.

### Questionable practices of correction for FPR under NHST

Univariate GLM is inefficient in handling neuroimaging data. It may work reasonably well if the following two conditions can be met: 1) no multiple testing, and 2) high signal-to-noise ratio (strong effect and high precision measurement) as illustrated in the lower triangular part of the table or right side of the curves in Fig. 4 (Appendix B). However, neither of the two conditions is likely satisfied with typical neuroimaging data. Due to the stringent requirements of correction for multiple testing across thousands of resolution elements in neuroimaging, a daunting challenge facing the community is the power inefficiency or high type II errors under NHST. Even if prior information is available as to which ROIs are potentially involved in a study, an ROI-based univariate GLM would still be obliged to share the burden of correction for multiplicity equally and agnostically to any such knowledge. The modeling approach usually does not have the luxury to survive the penalty, as shown with our experimental data in the table of Fig. 1, unless only a few ROIs are included in the analysis. Furthermore, with many low-hanging fruits with relatively strong signal strength (e.g., 0.5% signal change or above) having been largely plucked, the typical effect under investigation nowadays is usually subtle and likely small (e.g., in the neighborhood of 0.1% signal change). Compounded with the presence of substantial amount of noise and suboptimal modeling (e.g., ignoring the HDR shape subtleties), detection power is usually low. It might be counterintuitive, but one should realize that the noisier or more variable the data, the less one should be confident about any inferences based on statistical significance, as illustrated with the type S and type M errors in Figure 5 (Appendix B). With fixed threshold correction approaches, small regions are hard to funnel through the FPR criterion even if their effect magnitude is the same as or even higher than those larger regions. Even for approaches that take into consideration both spatial extent and effect magnitude (TFCE, ETAC), small regions remain disadvantaged when their effect magnitude is at the same level as their larger counterparts.

**Figure 4:**
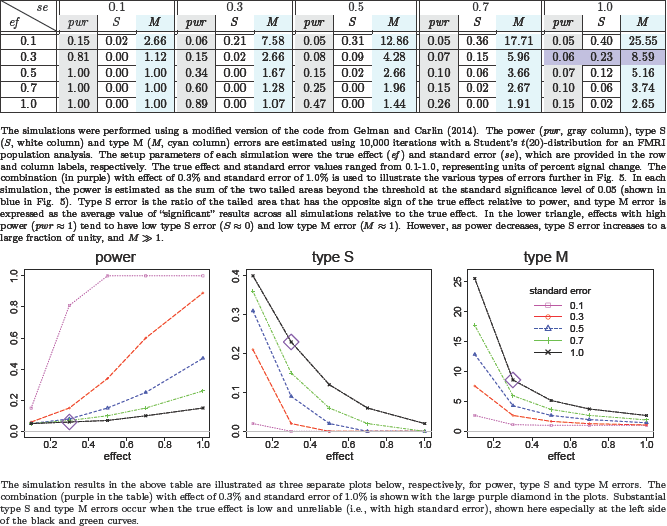
Power, type S and type M errors estimated from simulations

**Figure 5:**
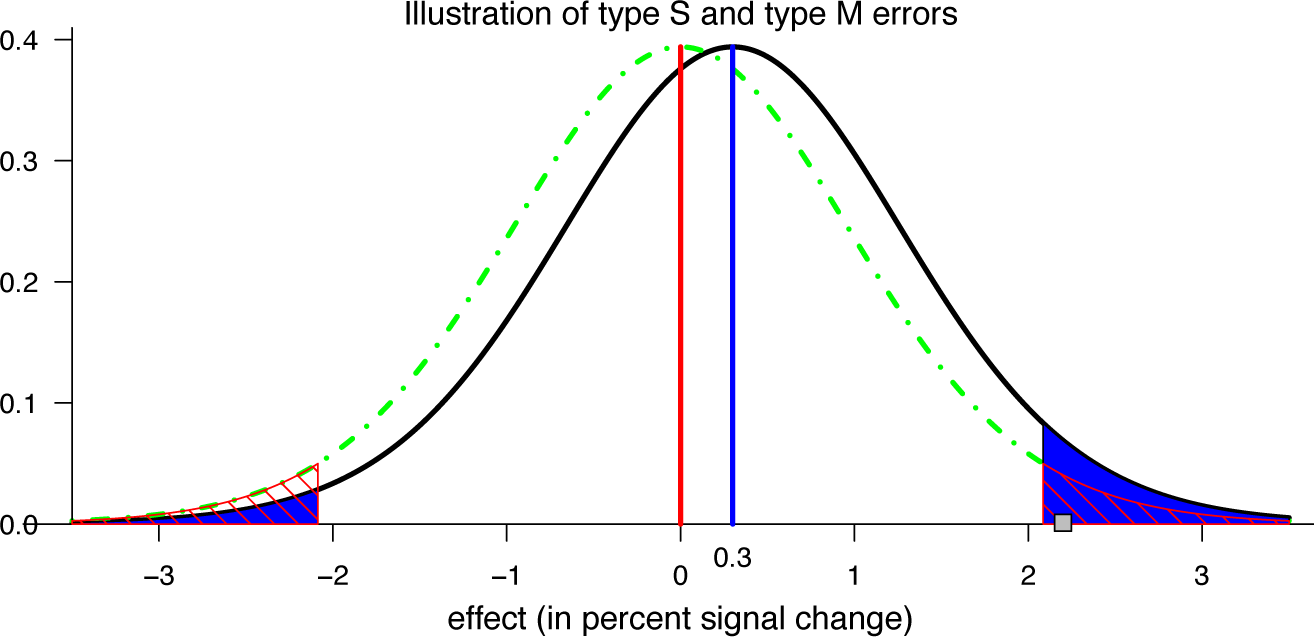
Illustration of the concept and interpretation for power, type I, type S and type M errors (Gelman, 2015).
Suppose that there is a hypothetical Student’s t(20)-distribution (black curve) for a true effect (blue vertical line of 0.3 and a corresponding standard error of 1.0 percent signal change, a scenario highlighted in purple in Fig. 4. Under the null hypothesis (red vertical line and dot-dashed green curve), two-tailed testing with a type I error rate of 0.05 leads to having thresholds at ±2.086; FPR = 0.05 corresponds to the null distribution’s total area beyond these two critical values (marked with red diagonal lines). The power is the total area of the t(20)-distribution for the true effect (black curve) beyond these thresholds, which is 0.06 (shaded in blue). The type S error is the ratio of the blue area in the true effect distribution’s left tail beyond the threshold of −2.086 to the area in both tails, which is 23% here (i.e., the ratio of the “significant” area in the wrong-signed tail to that of the total “significant” area). If a random draw from the t(20)-distribution under the true effect happens to be 2.2 (small gray square), it would be identified as statistically significant at the 0.05 level, and the resulting type M error would quantify the magnification of the estimated effect size as 2.2/0.3 ≈ 7.33, which is much larger than unity

Furthermore, dichotomous thinking and decision-making under NHST are usually not fully compatible with the underlying mechanism under investigation. Current knowledge regarding brain activations has not reached a point where one can make accurate dichotomous claims as to whether a specific brain region under a condition is activated or not; lack of underlying “ground truth” has made it difficult to validate any but the most basic models. The same issue can be raised about the binary decision as to whether the response difference under two conditions is either the same or different. Therefore, a pragmatic mission is to detect activated regions in terms of practical, instead of statistical, significance. The conventional NHST runs against the idealistically null point **H*_0_*, and declares a region having no effect based on statistical significance with a safeguard set for type I error. When the power is low, not only reproducibility will suffer, but also the chance of having an incorrect sign for a statistically significant effect be substantial (Fig. 4 and Fig. 5). Only when the power reaches 30% or above does the type S error rate become low. Publication bias due to the thresholding funnel contributes to type S and type M errors as well. The sharp thresholding imposed by the widely adopted NHST strategy uses a single threshold through which a high-dimension dataset is funneled. An ongoing debate has been simmering for a few decades regarding the legitimacy of NHST, ranging from cautionary warning against misuses of NHST (Wasserstein and Lazar, 2016), to tightening the significance level from 0.05 to 0.005 (Benjamin et al., 2017), to totally abandoning NHST as a gatekeeper (McShare et al., 2017; Amrhein and Greenland, 2017). The poor controllability of type S and type M errors is tied up with widespread problems across many fields. It is not a common practice nor a requirement in neuroimaging to report the effect estimates; therefore, power analysis for a brain region under a task or condition is largely obscure and unavailable, let alone the assessment of type S and type M errors.

Lastly, reproducibility may deteriorate through inefficient modeling and dichotomous inferences. Relating the discussion to the neuroimaging context, the overall or global FPR is the probability of having data as extreme as or more extreme than the current result, under the assumption that the result was produced by some “random number generator,” which is built into algorithms such as Monte Carlo simulations, random field theory, and randomly shuffled data as pseudo-noise in permutations. It boils down to the question: are the data truly pure noise (even though spatially correlated to some extent) in most brain regions? Since controlling for FPR hinges on the null hypothesis of no effect, p-value itself is a random variable that is a nonlinear continuous function of the data at hand, therefore it has a sampling distribution (e.g., uniform(0,1) distribution if the null model is true). In other words, it might be under-appreciated that even identical experiments would not necessarily replicate an effect that is dichotomized in the first one as statistically significant (Lazzeroni et al., 2016). The common practice of reporting only the statistically significant brain regions and comparing to those nonsignificant regions based on the imprimatur of statistic- or p-threshold can be misleading: the difference between a highly significant region and a nonsignificant region could simply be explained by pure chance. The binary emphasis on statistical significance unavoidably leads to an investigator only focusing on the significant regions and diverting attention away from the nonsignificant ones. More importantly, the traditional whole brain analysis usually leads to selectively report surviving clusters conditional on statistical significance through dichotomous thresholding, potentially inducing type M errors, biasing estimates with exaggerated effect magnitude, as illustrated in Fig. 5. Rigor, stringency, and reproducibility are lifelines of science. We think that there is more room to improve the current practice of NHST and to avoid information waste and inefficiency. Because of low power in FMRI studies and the questionable practice of binarized decisions under NHST, a Bayesian approach combined with integrative modeling offers a platform to more accurately account for the data structure and to leverage the information across multiple levels.

### What Bayesian modeling offers

Bayesian inferences are usually more compatible with the research, not null, hypothesis. Almost all statistics consumers (including the authors of this paper) were *a priori* trained within the conventional NHST paradigm, and their mindsets are usually familiar with and entrenched within the concept and narratives of *p*-value, type I error, and dichotomous interpretations of results. Out of the past shadows cast by the theoretical and computation hurdles, as well as the image of subjectivity, Bayesian methods have gradually emerged into the light. One direct benefit of Bayesian inference, compared to NHST, is its concise and straightforward interpretation, as illustrated in Table 5. For instance, the conventional confidence interval weighs equally all possible values a parameter could take within the interval, regardless of how implausible some of them are; in contrast, the quantile interval under Bayesian framework is more subtly expressed through the corresponding posterior density (Fig. 2). Even though the NHST modeling strategy literally falsifies the straw man *H*_0_, the real intention is to confirm the alternative (or research) hypothesis through rejecting *H*_0_; in other words, the falsification of *H*_0_ is only considered an intermediate step under NHST, and the ultimate goal is the confirmation of the intended hypothesis. In contrast, under the Bayesian paradigm, the investigator’s hypothesis is directly placed under scrutiny through incorporating prior information, model checking and revision. Therefore, the Bayesian paradigm is more fundamentally aligned with the hypothetico-deductivism axis along the classic view of falsifiability or refutability by Karl Popper (Gelman and Shalizi, 2013).

In addition to interpretational convenience, Bayesian modeling is less vulnerable to the amount of data available and to the issue of multiple testing. Conventional statistics heavily relies on large sample size and asymptotic property; in contrast, Bayesian inferences bear a direct interpretation conditional on the data regardless of sample size. Practically speaking, should we fully “trust” the effect estimate at each ROI or voxel at its face value? Some may argue that the effect estimate from the typical neuroimaging individual or population analysis has the desirable property of unbiasedness, as asymptotically promised by central limit theory. However, in reality the asymptotic behavior requires a very large sample size, a luxury difficult for most neuroimaging studies to reach. As the determination of a reasonable sample size depends on signal strength, brain region, and noise level, the typical sample size in neuroimaging tends to get overstretched, creating a hotbed for a low power situation and potentially high type S and type M errors. Another fundamental issue with the conventional univariate modeling approach in neuroimaging is the two-step process: first, pretend that the voxels or nodes are independent with each other, and build a separate model for each spatial element; then, handle the multiple testing issue using spatial relatedness to only partially, not fully, recoup the efficiency loss. In addition to the conceptual novelty for those with little experience outside the NHST paradigm, BML simplifies the traditional two-step workflow with one integrative model, fully shunning the multiple testing issue.

Through integrative incorporation across ROIs, the BML approach renders conservative effect estimation in place of controlling FPR. Unlike the *p*-value under NHST, which represents the probability of obtaining the current data generated by an imaginary machinery (e.g., assuming that no effect exists), a posterior distribution for an effect under the Bayesian framework directly and explicitly shows the uncertainty of our current knowledge conditional on the current data and the prior model. From the Bayesian perspective, we should not put too much faith in point estimates. In fact, not only should we not fully trust the point estimates from GLM with no pooling, but also should we not rely too much on the centrality (median, mean) of the posterior distribution from a Bayesian model. Instead, the focus needs to be placed more on the uncertainty, which can be visualized through the posterior distributions or characterized by the quantile intervals of the posterior distribution if summarization is required. Specifically, BML, as demonstrated here with the ROI data, is often more conservative in the sense that it does not “trust” the original effect estimates as much as GLM, as shown in Fig. 1; additionally, in doing so, it fits the data more accurately than the ROI-based GLM (Table 4c and Fig. 3). Furthermore, the original multiple testing issue under the massively univariate platform is deflected within one unified model: it is the type S, not type I, errors that are considered crucial and controlled under BML. Even though the posterior inferences at the 95% quantile interval in our experimental data were similar to the statistically significant results at the 0.05 level under NHST, BML in general is more statistically conservative than univariate GLM under NHST, as shown with the examples in Gelman et al. (2012).

We reiterate that the major difference is the assumption about the brain regions: noninformitive flat prior for the conventional GLM versus the Gaussian assumption for BML. With a uniform prior, all values on the real axis are equally likely; therefore, no information is shared across regions under GLM. On the other hand, it is worth mentioning that the Gaussianity assumption for the priors including the likelihood under a Bayesian model is based on two considerations: one aspect is convention and pragmatism, and the other aspect is the fact that, per maximum entropy principle, the most conservative distribution is Gaussian if the data have a finite variance (McElreath, 2016). However, a Bayesian model tends to be less sensitive to the model (likelihood or prior for the data in Bayesian terminology); in other words, even though the true data-generating process is always unknown, a model is only a convenient framework or prior knowledge to start with the Bayesian updating process so that a Bayesian model is usually less vulnerable to assumption violations. In contrast, statistical inferences with conventional approaches heavily rely on the sampling distribution assumptions. Through adaptive regularization, BML achieves a goal to trade off poorer fit in sample for better inference and improved fit out of sample (McElreath, 2016); the amount of regularization is learned from the data through partial pooling that embodies the similarity assumption of effects among the brain regions. From the NHST perspective, BML can still commit type I errors, and its FPR could be higher under some circumstances than, for example, its GLM counterpart. Such type I errors may sound alarmingly serious; however, the situation is not as severe as its appearance for two reasons: 1) the concept of FPR and the associated model under NHST are framed with a null hypothesis, which is not considered pragmatically meaningful in the Bayesian perspective; and 2) in reality, inferences under BML most likely have the same directionality as the true effect because type S errors are well controlled across the board under BML (Gelman and Teulinckx, 2000). Just consider which of the following two scenarios is worse: (a) when power is low, the likelihood under the NHST to mistakenly infer that the BOLD response to easy condition is higher than difficult could be sizable (e.g., 30%), and (b) with the type S error rate controlled below, for example, 3.0%, the BML might exaggerate the magnitude difference between difficult and easy conditions by, for example, 2 times. While not celebrating the scenario (b), we expect that most researchers would view the scenario (a) as more problematic.

Prior assignment is an intrinsic component of Bayesian modeling. One somewhat controversial aspect of Bayesian modeling is the adoption of a prior for each hyperparameter, and that the prior is meshed with the data and gets updated into the posterior distribution. Some may consider that an Achilles’ heel of Bayesian modeling is its subjectivity with respect to prior selection. First of all, we would argue that, to start with, every model, Bayesian or non-Bayesian, is a prior or likelihood function in the sense that the analyst presumes a distribution for the data (e.g., the conventional GLMs (1), (2), and (3)). Secondly, weakly informative priors are even routinely adopted by conventional statistics in approaches such as penalized likelihood in ridge regression and LASSO. Priors are chosen, evaluated and revised just as any components and assumptions in the model. The inherent subjectivity of BML is no more than are model assumptions (e.g. Gaussianity) for convectional statistics as well as different opinions of analytical approaches, different outliers handling methods and processing steps in neuroimaging. Furthermore, priors are applied at the epistemological, not ontological level (McElreath, 2016); with no intention to get tangled in the epistemological roots or the philosophical debates of subjectivity versus objectivity, we simply divert the issue to the following suggestion (Gelman and Hennig, 2017): replacing the term “subjectivity” with “multiple perspectives and context dependence,” and, in lieu of “objectivity,” focusing on transparency, consensus, impartiality, and correspondence to observable reality. Therefore, our focus here is the pragmatic aspect of Bayesian modeling: with prior distributions, we can make inferences for each ROI under BML, which cannot be achieved under LME.

Prior selection in Bayesian modeling is usually well justified. Since noninformative priors are adopted for population effects, the only impact of prior information incorporated into BML comes from two aspects: the distributional assumptions about those entities such as subjects and ROIs, and the hyperpriors. However, the rationale for the Gaussian priors of entities is not more far-fetched than that for the Gaussianity of cross-subject distribution in the typical GLM adopted in neuroimaging group analysis. As for the hyperpiors, they basically play the role of regularization: when the amount of data is large, the weakly informative priors levy a negligible effect on the final inferences; on the other hand, when the data do not contain enough information for robust inferences, the weakly informative priors prevent the distributions from becoming unsupportively dispersive. Specifically, for simple models such as Student’s t-test and GLM, Bayesian approach renders similar inferences if noninformative priors are assumed. On the surface a noninformative prior does not “artificially” inject much “subjective” information into the model, and should be preferred. In other words, it might be considered a desirable property from the NHST viewpoint, since noninformative priors are independent of the data. Because of this “objectivity” property, one may insist that noninformative priors should be used all the time. Counterintuitively, a noninformative prior may become so informative that it could cause unbounded damage (Gelman et al., 2017). If we analyze the r ROIs individually as in the *r* GLMs (14), the point estimate for each effect **θ_j_** is considered stationary, and we would have to correct for multiple testing due to the fact that *r* separate models are fitted independently. Bonferroni correction would likely be too harsh, which is the major reason that ROI-based analysis is rarely adopted in neuroimaging except for effect verification or graphic visualization. On the other hand, the conventional approach with the *r* GLMs (14) is equivalent to the BML (17) by *a priori* assuming an improper flat prior for **θ_j_** with the cross-ROI variability τ^2^ = ∞ that is, each effect **θ_j_** can be any value with equal likelihood within (–∞,∞). In the case of BOLD response, it is not necessarily considered objective to adopt a noninformative priori such as uniform distribution when intentionally ignoring the prior knowledge. In fact, we do have the prior knowledge that the typical effect from a 3T scanner has the same scale and lies within, for example, (–4,4) in percent signal change; this commonality can be utilized to calibrate or regularize the noise, extreme values, or outliers due to pure chance or unaccounted-for confounding effects (Gelman et al., 2012), which is the rationale for our prior distribution assumption of Gaussianity for both subjects and ROIs. Even for an effect for a covariate (e.g., the associate between behavior and BOLD response), it would be far-fetched to assume that τ^2^ has the equal chance between, for example, 0 and 10^10^. Another example of information waste under NHST is the following. Negative or zero variance can occur in an ANOVA model while zero variance may show up in LME. Such occurrences are usually due to the full reliance on the data or a parameter boundary, and such direct estimates are barely meaningful: an estimate of cross-subject variability λ^2^ = 0 in (17) indicates that all subjects have absolutely identical effects. However, a Bayesian inference is a tug of war between data and priors, and therefore negative or zero variance inferences would not occur because those scenarios from the data are regularized by the priors, as previously shown in ICC computations that are regularized by a Gamma prior for the variance components (Chen et al., 2017c).

In typical neuroimaging data with reasonable number of subjects and moderate number of ROIs, the weakly informative priors for scaling parameters usually play a nudging role. In general, when there is enough data, weakly informative priors are usually drowned out by the information from the data; on the other hand, when data are meager (e.g., with small or moderate sample size), such priors can play the role of regularization (Gelman et al., 2017) so that smoother and more stable inferences could be achieved than would be obtained with a flat prior. In addition, a weakly informative prior for BML allows us to make reasonable inferences at the region level while model quality can be examined through tools such as posterior predictive check and LOO cross-validation. Therefore, if we do not want to waste such prior knowledge for an effect bounded within a range in the brain, the commonality shared by all the brain regions can be incorporated into the model through a weakly informative prior and the calibration of partial pooling among the ROIs, thus eliding the step of correcting for multiple testing under NHST.

To summarize, we recommend that three types of priors be adopted for ROI-based BML: 1) Gaussian distribution for the response variable (or input data) and effects at the entity level such as ROIs and subjects, 2) uninformative prior for the effects (e.g., intercept and slopes) at the population level, and 3) weekly informative priors for scaling parameters (e.g., variances). Such a prior setting should be able to handle the typical BML modeling in neuroimaging unless the amount of data is overly meager (e.g., a few ROIs or subjects only). With the typical neuroimaging dataset, our prior recommendation would have negligible impact on the population effect estimates as shown in the comparisons between LME and BML (Table 4). However, it is worth noting that the effects of interest here are not those at the population level under the LME and BML framework, but rather those effects at the entity (i.e. ROI) level in the current context.

Progress in Bayesian computations is paving the way for more advanced modeling opportunities. Bayesian algorithms have traditionally been burdened with meticulous and time consuming computations, making their adoption for wide applications difficult and impractical for multilevel models even with a dataset of small or moderate size. However, the situation has been substantially ameliorated by the availability of multiple software tools such as Stan, and the rapid development in Stan over the past few years has promoted the wide adoption of Bayesian modeling. In particular, Stan adopts static HMC Samplers and its extension, NUTS, and it renders less autocorrelated and more effective draws for the posterior distributions, achieving quicker convergence than the traditional Bayesian algorithms such as Metropolis-Hastings, Gibbs sampling, and so on. With faster convergence and high efficiency, it is now feasible to perform full Bayesian inferences for BML with datasets of moderate size.

### Advantages of ROI-based BML

Bayesian modeling has long been adopted in neuroimaging at the voxel or node level (e.g., Woolrich et al., 2004; Penny et al., 2005; Westfall et al., 2017; Eklund et al., 2017; Mejia et al., 2017); nevertheless, correction for FWE would still have to be imposed as part of the model or as an extra step. In the current context, we formulate the data generation mechanism for each dataset through a progressive triplet of models on a set of ROIs: GLM → LME → BML. The strength of multilevel modeling lies in its capability of stratifying the data in a hierarchical or multilevel structure so that complex dependency or correlation structures can be properly accounted for coherently within a single model. Specifically applicable in neuroimaging is a crossed or factorial layout between the list of ROIs and subjects as shown in the LME equation (4) and its Bayesian counterpart (5). Our adoption of BML, as illustrated with the demonstrative data analysis, indicates that BML holds some promise for neuroimaging and offers the following advantages over traditional approaches:

1) Compared to the conventional GLM, BML achieves a better model performance and higher predictive accuracy through partial pooling, a trade-off between underfitting with complete pooling and overfitting with no pooling. Specifically, BML is assessed with data through adaptive regularization with nudges from the prior information: it learns from the data and borrows information across ROIs to improve the quality of individual estimates and posterior distributions with the assumption of similarity among the regions.
2) Instead of separately correcting for multiple testing, BML incorporates multiple testing as part of the model by assigning a prior distribution among the ROIs (i.e., treating ROIs as random effects under the LME paradigm). In doing so, multiple testing is handled under the scaffold of the multilevel data structure by conservatively shrinking the original effect toward the center; that is, instead of leveraging cluster size or signal strength, BML leverages the commonality among ROIs.
3) BML may achieve higher spatial specificity through efficient modeling. A statistically identified cluster through a whole brain analysis is not necessarily anatomically or functionally meaningful. In other words, a statistically identified cluster is not always aligned well with a brain region for diverse reasons such as “bleeding” effect due to contiguity among regions, and suboptimal alignment to the template space, as well as spatial blurring. In fact, a cluster may overlap multiple brain regions or subregions. In contrast, as long as a region can be *a priori* defined, its statistical inference under BML is assessed by its signal strength relative to its peers, not by its spatial extent, providing an alternative to the whole brain analysis with more accurate spatial specificity.
4) BML offers a flexible approach to dealing with double sidedness at the ROI level. When prior information about the directionality of an effect is available on some, but not all, regions (e.g., from previous studies in the literature), one may face the issue of performing two one-tailed t-tests at the same time in a blindfold fashion due to the limitation of the massively univariate approach. The ROI-based approach disentangles the complexity since the posterior inference for each ROI can be made separately.
5) Model validation is a crucial facet of Bayesian framework. It may be trite to cite the famous quote of “all models are wrong” by George E. P. Box. However, the reality in neoroimaging is that model quality check is substantially lacking. When prompted, one may recognize the potential problems and pitfalls of a model. Nevertheless, statistical analysis is typically conducted as mechanical operations on assembly lines; when discussing and reporting results from the model, the investigator tends to treat the model as if it were always true and then discusses statistical inferences without realizing the implications or ramifications of the model that fits poorly or even conflicts with data. Building, comparing, tuning and improving models is a daunting task with whole brain data due to the high computational cost and visualization inconvenience. In contrast, model quality check is an intrinsic part of Bayesian modeling process. The performance of each model and the room for improvement can be directly examined through graphical display as shown in Fig. 3.
6) A full results reporting is possible for all ROIs under BML. The conventional NHST focuses on the point estimate of an effect supported with statistical evidence in the form of a p-value. In the same vein, typically the results from the whole brain analysis are displayed with sharp-thresholded maps or tables that only show the surviving clusters with statistic- or p-values. In contrast, as the focus under the Bayesian framework is on the predictive distribution, not the point estimate, of an effect, the totality of BML results can be summarized in a table as shown in Figures 1 and 2, listing the predictive intervals in various quantiles (e.g., 50%, 75%, and 95%), a luxury that whole brain analysis cannot provide. Such totality pits against the backdrop in which the effect estimates are usually not reported in the whole brain analysis (Chen et al., 2017b). In contrast, BML modeling at the ROI level directly allows the investigator to present the effect estimate. More importantly, BML substantiates the reporting advantage not only because of modeling at the ROI level, but also due to the fact that the uncertainty associated with each effect estimate can demonstrated in a much richer fashion (e.g., explicit revealing the spread or skewness of the posterior distribution) through the posterior density distribution (Fig. 2) than the conventional confidence interval (1) that is flat and inconvenient to interpret.
7) To some extent, the ROI-based BML approach can alleviate the arbitrariness involved in the thresholding with the current FPR correction practices. Even though BML allows the investigator to present the whole results for all regions, for example, in a table format, we do recognize that the investigator may prefer to focus the discussion on some regions with strong posterior evidence. In general, with all effects reported in totality, regardless of their statistical evidence, the decision of choosing which effects to discuss in a paper should be based on cost, benefit, and probabilities of all results (Gelman et al., 2014). Specifically for neuroimaging data analysis, the decision still does not have to be solely from the posterior distribution; instead, we suggest that the decision be hinged on the statistical evidence from the current data, combined with prior information from previous studies. For example, one may still choose the 95% quantile interval as an equivalent benchmark to the conventional p-value of 0.05 when reporting the BML results. However, those effects with, say, 90% quantile intervals excluding 0 can still be discussed with a careful and transparent description, which can be used as a reference for future studies to validate or refute; or, such effects can be reported if they have been shown in previous studies. Moreover, rather than a cherry-picking approach on reporting and discussing statistically significant clusters in whole brain analysis^6^, we recommend a principled approach in results reporting in which the ROI-based results be reported in totality with a summary as shown in Figures 1 and 2 and be discussed through transparency and soft, instead of sharp, thresholding. We believe that such a soft thresholding strategy is more healthy and wastes less information for a study that goes through a strenuous pipeline of experimental design, data collection, and analysis.

### Limitations of ROI-based BML and future directions

ROIs can be specified through several ways depending on the specific study or information available regarding the relevant regions. For example, one can find potential regions involved in a task or condition including resting state from the literature. Such regions are typically reported as the coordinates of a “peak” voxel (usually highest statistic value within a cluster), from which each region could be defined by centering a ball with a radius of, e.g., 6 mm in the brain volume (or by projecting an area on the surface). Regions can also be located through (typically coordinate-based) meta analysis with databases such as NeuroSynth (http://www.neurosynth.org) and BrainMap (http://www.brainmap.org), with tools such as brain_matrix (https://github.com/fredcallaway/brain_matrix), GingerALE (http://brainmap.org/ale), Sleuth (http://brainmap.org/sleuth), and Scribe (http://www.brainmap.org/scribe) that are associated with the database BrainMap. Anatomical atlases (e.g., http://surfer.nmr.mgh.harvard.edu, href="http://www.med.harvard.edu/aanlib) and functional parcellations (e.g., Schaefer et al., 2017) are another source of region definition. As a different strategy, by recruiting enough subjects, one could use half of the subjects to define ROIs, and the other half to perform ROI-based analysis; similarly, one could scan the same set of subjects longer, use the first portion of the data to define ROIs, and the rest to perform ROI-based analysis.

The limitations of the ROI-based BML are as follows.

1) Just as the FWE correction on the massively univariate modeling results is sensitive to the size of the full domain in which it is levied (whole brain, gray matter, or a user-defined volume), so the results from BML will depend to some extent on the number of ROIs (or which) ones included. For a specific ROI *j*, changing the composition among the rest of ROIs (e.g., adding an extra ROI or replacing one ROI with another) may result in a different prior distribution (e.g, **θ_j_** ~ N(μ, τ^2^) in BML (5)) and a different posterior distribution for *θ_j_*. However, it merits noting that the regions should not be arbitrarily chosen but rather selected from the current knowledge and relevancy of the involving effect under investigation.
2) ROI data extraction involves averaging among voxels within the region. Averaging, as a spatial smoothing or low-pass filtering process, condenses, reduces or dilutes the information among the voxels (e.g., 30) within the region to one number, and loses any finer spatial structure within the ROI. In addition, the variability of extracted values across subjects and across ROIs could be different from the variability at the voxel level.
3) ROI-based analysis is conditional on the availability and quality of the ROI definition. One challenge facing ROI definition is the inconsistency in the literature due to inaccuracies across different coordinate/template systems and publication bias. In addition, some extent of arbitrariness is embedded in ROI definition; for example, a uniform adoption of a fixed radius may not work well due to the heterogeneity of brain region sizes. When not all regions or subregions currently can be accurately defined, or when no prior information is available to choose a region in the first place, the ROI-based approach may miss any potential regions if they are not included in the model.
4) The exchangeability requirement of BML assumes that no differential information is available across the ROIs in the model. Under some circumstances, ROIs can be expected to share some information and not fully independent, especially when they are anatomically contiguous or more functionally related than the other ROIs (e.g., homologous regions in opposite hemisphere). Ignoring spatial correlations, if substantially present, may lead to underestimated variability and inflated inferences. In the future we will explore the possibility of accounting for such a spatial correlation structure. A related question is whether we can apply the BML strategy to the whole brain analysis (e.g., shrinking the effects among voxels). Although tempting, such an extension faces serious issues, such as daunting computational cost, serious violation of the exchangeability assumption, and loss of spatial specificity. Despite these limitations, we believe that BML holds its unique promising potentials and advantages over the conventional approaches, and we hope that it will serve as a catalyst for a wider modeling landscape in neuroimaing. Admittedly, as all models are idealized statistical representations, our BML work presented here is only an incremental step in neuroimaging; besides multiplicity, NIHST pitfalls and inefficient modeling, there remain daunting challenges such as linearity assumption, temporal correlation (Olszowy et al., 2017) and the inaccuracy of presumed hemodynamic response modeling in FMRI data analysis.

### Conclusion

The prevalent adoption of dichotomous decision making under NHST runs against the continuous nature of most quantities under investigation, including neurological responses, which has been demonstrated to be problematic through type S and type M errors when the signal-to-noise ratio is low. The conventional correction for FWE in neuroimaging data analysis is viewed as a “desirable” standard procedure for whole brain analysis because the criterion is a pivotal component of NHST. However, it is physiologically unfeasible to claim that there is absolutely no effect in most brain regions; therefore, we argue that setting the stage only to fight the straw man of no effect anywhere is not necessarily a powerful nor efficient inference strategy. Inference power is further comprimised by FWE correction due to the inefficiency involved in the massively univariate modeling approach. As BOLD responses in the brain share the same scale and range, the ROI-based BML approach proposed here allows the investigator to borrow strength and effectively regularize the distribution among the regions, and it can simultaneously achieve meaningful spatial specificity and detection efficiency. In addition, it can provide increasing transparency on model building, quality control, and detailed results reporting, and offers a promising approach to addressing two multiplicity issues: multiple testing and double sidedness.

## Acknowledgments

The research and writing of the paper were supported (GC, PAT, and RWC) by the NIMH and NINDS Intramural Research Programs (ZICMH002888) of the NIH/HHS, USA, and by the NIH grant R01HD079518A to TR and ER. Much of the modeling work here was inspired from Andrew Gelman’s blog. We are indebted to Paul-Christian Bürkner and the Stan development team members Ben Goodrich, Daniel Simpson, Jonah Sol Gabry, Bob Carpenter, and Michael Betancourt for their help and technical support. The simulations were performed in R and the figures were generated with the R package ggplot2 (Wickham, 2009).

## Appendix A. Pitfalls of NHST

i. It is a common mistake by investigators and even statistical analysts to misinterpret the conditional probability under NHST as the posterior probability of the truth of the null hypothesis (or the probability of the null event conditional on the current data at hand) even though fundamentally *P(data* | **H*_0_)* ≠ *P(*H*_0_* | *data).*
ii. One may conflate statistical significance with practical significance, and subsequently treat the failure to reach statistical significance as the nonexistence of any meaningful effect. Even though the absence of evidence is not an evidence of absence, it is common to read discussions in scientific literature wherein the authors implicitly (or even explicitly) treat statistically non-significant effects as if they were zero.
iii. Statistic- or p-values cannot easily be compared: the difference between a statistically significant effect and another effect that fails to pass the significance level does not necessarily itself reach statistical significance.
iv. How should the investigator handle the demarcation, due to sharp thresholding, between one effect with p = 0.05 (or a surviving cluster cutoff of 54 voxels) and another with p = 0.051 (or a cluster size of 53 voxels)^7^?
v. The focus on statistic- or p-value seems to, in practice, lead to the preponderance of reporting only statistical, instead of effect, maps in neuroimaging, losing an effective safeguard that could have filtered out potentially spurious results (Chen et al., 2017b).
vi. One may mistakenly gain more confidence in a statistically significant result (e.g., high statistic value) in the context of data with relatively heavy noise or with a small sample size (e.g., leading to statement such as “despite the small sample size” or “despite the limited statistical power”). In fact, using statistical significance as a screener can lead researchers to make a wrong assessment about the sign of an effect or drastically overestimate the magnitude of an effect.
vii. While the conceptual classifications of false positives and false negatives make sense in a system of discrete nature (e.g., juror decision on *H*_0_: the suspect is innocent), what are the consequences when we adopt a mechanical dichotomous approach to assessing a quantity of continuous, instead of discrete, nature?
viii. It is usually under-appreciated that the p-value, as a function of data, is a random variable, and thus itself has a sampling distribution. In other words, p-values from experiments with identical designs can differ substantially, and statistically significant results may not necessarily be replicated (Lazzeroni et al., 2016).

## Appendix B. Type S and type M errors

We discuss two types of error that are not often discussed in neuroimaging: type S and type M. These two types of error cannot be directly captured by the FPR concept and may become severe when the effect is small relative to the noise, which is usually the situation in BOLD neuroimaging data. In the NHST formulation, we formulate a null hypothesis *H*_0_ (e.g., the effect of an easy task E is identical to a difficult one D), and then commit a type I (or false positive) error if wrongly rejecting *H*_0_ (e.g., the effect of easy is judged to be statistically significantly different from difficult when actually their effects are the same); in contrast, we make a type II (or false negative) error when accepting *H*_0_ when *H*_0_ is in fact false (e.g., the effect of easy is assessed to be not statistically significant from difficult even though their effects do differ). These are the dichotomous errors associated with NHST, and the counterbalance between these two types of error are the underpinnings of typical experimental design as well results reporting.

However, we could think about other ways of framing errors when making a statistical assessment (e.g., the easy case elicits a stronger BOLD response at some region than the difficult case) conditional on the current data. We are exposed to a risk that our decision is contrary to the truth (e.g., the BOLD response to the easy condition is actually lower than to the difficult condition). Such a risk is gauged as a type S (for “sign”) error when we incorrectly identify the sign of the effect; its values range from 0 (no chance of error) to 1 (full chance of error). Similarly, we make a type M (for “magnitude”) error when estimating the effect as small in magnitude if it is actually large, or when claiming that the effect is large in magnitude if it is in fact small (e.g., saying that the easy condition produces a *much* large response than the difficult one when actually the difference is tiny); its values range across the positive real numbers: [0, 1) correspond to underestimation of effect magnitude, 1 describes correct estimation, and (1, ∞^+^) mean overestimation. The two error types are illustrated in Fig. 5 for inferences made under NHST. In the neuroimaging realm, effect magnitude is certainly a property of interest, therefore the corresponding type S and type M errors would be of research interest.

Geometrically speaking, if the null hypothesis **H*_0_* can be conceptualized as the point at zero, NHST aims at the real space R excluding zero with a pivot at the point of zero (e.g., *D — E* = 0); in contrast, type S error gauges the relative chance that a result is assessed on the wrong side of the distribution between the two half spaces of R (e.g., D — E > 0 or D — E < 0), and type M error gauges the relative magnitude of differences along segments of R+ (e.g., the ratio of *measured* to *actual* effect is >> 1 or << 1). Thus, we characterize type I and type II errors as “point-wise” errors, driven by judging the equality, and describe type S and type M errors as “direction-wise,” driven by the focus of inequality or directionality.

One direct application of type M error is that publication bias can lead to type M errors, as large effect estimates are more likely to filter through the dichotomous decisions in statistical inference and reviewing process. Using the type S and type M error concepts, it might be surprising for those who encounter these two error types for the first time to realize that, when the data are highly variable or noisy, or when the sample size is small with a relatively low power (e.g., 0.06), a statistically significant result at the 0.05 level is quite likely to have an incorrect sign - with a type S error rate of 24% or even higher (Gelman and Carlin, 2014). In addition, such a statistically significant result would have a type M error with its effect estimate much larger (e.g., 9 times higher) than the true value. Put it another way, if the real effect is small and sampling variance is large, then a dataset that reaches statistical significance must have an exaggerated effect estimate and the sign of the effect estimate is likely to be incorrect. Due to the ramifications of type M errors and publication filtering, an effect size from the literature could be exaggerated to some extent, seriously calling into question the usefulness of power analysis under NHST in determining sample size or power, which might explain the dramatic contrast between the common practice of power analysis as a requirement for grant applications and the reproducibility crisis across various fields. Fundamentally, power analysis inherits the same problem with NHST: a narrow emphasis on statistical significance is placed as a primary focus (Gelman and Carlin, 2013).

The typical effect magnitude in BOLD FMRI at 3 Tesla is usually small, less than 1% signal change in most brain regions except for areas such as motor and primary sensory cortex. Such a weak signal can be largely submerged by the overwhelming noise and distortion embedded in the FMRI data. The low power for detection of typical FMRI data analyses in typical datasets is further compounded by the modeling challenges in accurately capturing the effect. For example, even though large number of physiological confounding effects are embedded in the data, it is still difficult to properly incorporate the physiological “noises” (cardiac and respirary effects) in the model. Moreover, habituation, saturation, or attenuation across trials or within each block are usually not considered, and such fluctuations relative to the average effect would be treated as noise or fixed-instead of random-effects (Westfall et al., 2017). There are also strong indications that a large portion of BOLD activations are usually unidentified at the individual subject level due to the lack of power ( Gonzalez-Castillo et al., 2012). Because of these factors, the variance due to poor modeling overwhelms all other sources (e.g., across trials, runs, and sessions) in the total data variance (Gonzalez-Castillo et al., 2016); that is, the majority (e.g., 60-80%) of the total variance in the data is not properly accounted for in statistical models.

## Appendix C. Multiplicity in neuroimaging

In general, we can classify four types of multiplicity issues that commonly occur in neuroimaging data analysis.

A) Multiple testing. The first and major multiplicity arises when the same design (or model) matrix is applied multiple times to different values of the response or outcome variable, such as the effect estimates at the voxels within the brain. As the conventional voxel-wise neuroimaging data analysis is performed with a massively univariate approach, there are as many models as the number of voxels, which is the source of the major multiplicity issue: multiple testing. Those models can be, for instance, Student’s t-tests, AN(C)OVA, univariate or multivariate GLM, LME or Bayesian model. Regardless of the specific model, all the voxels share the same design matrix, but have different response variable values on the left-hand side of the equation. With human brain size on the order of 10^6^ mm^3^, the number of voxels may range from 20,000 to 150,000 depending on the voxel dimensions. Each extra voxel adds an extra model and leads to incrementally mounting odds of pure chance or “statistically significant outcomes,” presenting the challenge to account for the occurrence of mounting family-wise error (FWE), while effectively holding the overall false positive rate (FPR) at a nominal level (e.g., 0.05). In the same vein, surface-based analysis is performed with 30,000 to 50,000 nodes (Saad et al., 2004), sharing a similar multiple testing issue with its volume-based counterpart. Sometimes the investigator performs analyses at smaller number of regions of interest (ROIs), perhaps of order 100, but even here adjustment is still required for the multiple testing issue (though it is often not made).
B) *Double sidedness.* Another occurrence of multiplicity is the widespread adoption of two separate one-sided (or one-tailed) tests in neuroimaging. For instance, the comparison between the two conditions of “easy” and “difficult” are usually analyzed twice for the whole brain: one showing whether the easy effect is higher than difficult, and the other for the possibility of the difficult effect being higher than easy. One-sided testing for one direction would be justified if prior knowledge is available regarding the sign of the test for a particular brain region. When no prior information is available for all regions in the brain, one cannot simply finesse two separate one-sided tests in place of one two-sided test, and a double sidedness practice warrants a Bonferroni correction because the two tails are independent with respect to each other (and each one-sided test is more liberal than a two-sided test at the same significance level). However, simultaneously testing both tails in tandem for whole brain analysis without correction is widely used without clear justification, and this forms a source of multiplicity issue that needs proper accounting.
C) *Multiple comparisons.* It rarely occurs that only one statistical test is carried out in a specific neuroimaging study, such as a single one-sample t-test. Therefore, a third source of multiplicity is directly related to the popular term, multiple comparisons, which occur when multiple tests are conducted under one model. For example, an investigator that designs an emotion experiment with three conditions (easy, difficult, and moderate) may perform several separate tests: comparing each of the three conditions to baseline, making three pairwise comparisons, or testing a linear combination of the three conditions (such as the average of easy and difficult versus moderate). However, neuroimaging publications seldom consider corrections for such separate tests.
D) *Multiple paths.* The fourth multiplicity issue to affect outcome interpretation arises from the number of potential preprocessing, data dredging and analytical pipelines (Carp, 2012). For instance, all common steps have a choice of procedures: outlier handling (despiking, censoring), slice timing correction (yes/no, various interpolations), head motion correction (different interpolations), different alignment methods from EPI to anatomical data plus upsampling (1 to 4 mm), different alignment methods to different standard spaces (Talairach and MNI variants), spatial smoothing (3 to 10 mm), data scaling (voxel-wise, global or grand mean), confounding effects (slow drift modeling with polynomials, high pass filtering, head motion parameters), hemodynamic response modeling (different presumed functions and multiple basis functions), serial correlation modeling (whole brain, tissue-based, voxel-wise AR or ARMA), and population modeling (univariate or multivariate GLM, treating sex as a covariate of no interest (thus no interactions with other variables) or as a typical factor (plus potential interactions with other variables)). Each choice represents a “branching point” that could have a quantitative change to the final effect estimate and inference. Conservatively assuming three options at each step here would yield totally 3^10^ = 59,049 possible paths, commonly referred to as researcher degrees of freedom (Simmons et al., 2011). The impact of the choice at each individual step for this abbreviated list might be negligible, moderate, or substantial. For example, different serial correlation models may lead to substantially different effect estimate reliability (Olszowy et al., 2017); the estimate for spatial correlation of the noise could be sensitive to the voxel size to which the original data were upsampled (Mueller et al., 2017; Cox and Taylor, 2017), which may lead to different cluster thresholds and poor control to the intended FPR in correcting for multiplicity. Therefore, the cumulative effect across all these multilevel branching points could be a large divergence between any two paths for the final results. A multiverse analysis (Steegen et al., 2016) has been suggested for such situations of having a “garden of forking paths” (Gelman and Loken, 2013), but this seems highly impractical for neuroimaging data. Even when one specific analytical path is chosen by the investigator, it remains possible to invoke potential or implicit multiplicity in the sense that the details of the analytical steps such as data sanitation are conditional on the data (Gelman and Loken, 2013). The final interpretation of significance typically ignores the number of choices or the potential branchings that may affect the final outcome, even though it would be more preferable to have the statistical significance independent of these preprocessing steps.

## Appendix D. Bayesian modeling for one-way random-effects ANOVA

Here we discuss a classical framework, a hierarchical or multilevel model for a one-way random-effects ANOVA, and use it as a building block to expand to a Bayesian framework for neuroimaging group analysis. In evaluating this model, the controllability of inference errors will be focused on type S errors instead of the traditional FPR. Suppose that there are r measured entities (e.g., ROIs), with entity *j* measuring the effect *θ_j_* from *n_j_* independent Gaussian-distributed data points *yij*, each of which represents a sample (e.g., trial), i =1, 2, *…,nj*. The conventional statistical approach formulates r separate models,

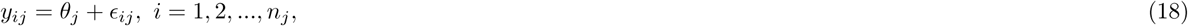

where *∊_i_j* is the residual for the *j*th entity and is assumed to be Gaussian N(0, σ^2^), *j =* 1, 2, *…,r.* Depending on whether the sampling variance σ^2^ is known or not, each effect can be assessed through its sample mean 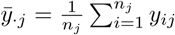 relative to the corresponding variance 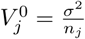, resulting in a Z- or t-test.

By combining the data from the r entities and further decomposing the effect **θ_j_** into an overall effect *b*_0_ across the *r* entities and the deviation ξ_j_ of the *j*th entity from the overall effect (i.e., **θ_j_* = *b*_0_ ξ*, *j* = 1, 2,…, r), we have a conventional one-way random-effects ANOVA,

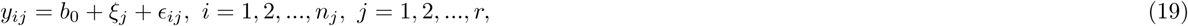

where *b*_0_ is conceptualized as a fixed-effects parameter, ξ_j_ codes the random fluctuation of the *j*th entity from the overall mean *b*_0_, with the assumption of ξ_j_ ~ N(0,r^2^), and the residual ∊_ij_ follows a Gaussian distribution N(0, σ^2^). The classical one-way random-effects ANOVA model (19) is typically formulated to examine the null hypothesis,

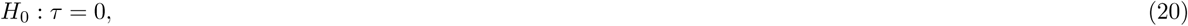

with an F-statistic, which is constructed as the ratio of the *between* mean sums of squares and the *within* mean sums of squares. An application of this ANOVA model (19) to neuroimaging is to compute the intraclass correlation ICC(1,1) as 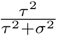 when the measuring entities are exchangeable (e.g., families with identical twins; Chen et al., 2017c).

Whenever multiple values (e.g., two effect estimates from two scanning sessions) from each measuring unit (e.g., subject or family) are correlated (e.g., the levels of a within-subject or repeated-measures factor), the data can be formulated using a linear mixed-effects (LME) model, sometimes referred to as a multilevel or hierarchical model. One natural ANOVA extension is simply to treat the model conceptually as LME, without the need of reformulating the model equation (19). However, LME can only provide point estimates for the overall effect *b*_0_, cross-region variance τ^2^ and the data variance σ^2^; that is, the LME (19) cannot directly provide any information regarding the individual *ξ_j_* or **θ_j_** values because of over-fitting due to the fact that the number of data points is less than the number of parameters that need to be estimated.

Our interest here is neither to assess the variability τ^2^ nor to calculate ICC, but instead to make statistical inferences about the individual effects *θ_j_*. Nevertheless, the conventional NHST (20) may shed some light on potential strategies (Gelman et al., 2014) for *θ_j_*. If the deviations ξ_j_ are relatively small compared to the overall mean *b*_0_, then the corresponding F-statistic value will be small as well, leading to the decision of not rejecting the null hypothesis (20) at a reasonable, predetermined significance level (e.g., 0.05); in that case, we can estimate the equal individual effects *θ_j_* using the overall weighted mean 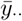. through full pooling with all the data,

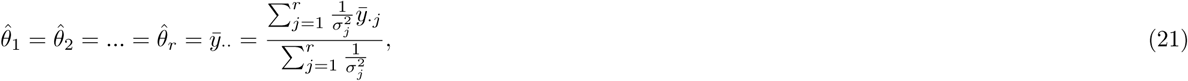

where 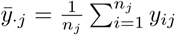 and 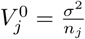 are the sampling mean and variance for the *j*th measuring entity, and the subscript dot (•) notation indicates the (weighted) mean across the corresponding index(es). On the other hand, if the deviations ξ_j_ are relatively large, so is the associated F-statistic value, leading to the decision of rejecting the null hypothesis (20); similarly, we can reasonably estimate *θ_j_* with no pooling across the r entities; that is, each *θ_j_* is estimated using the *j*th

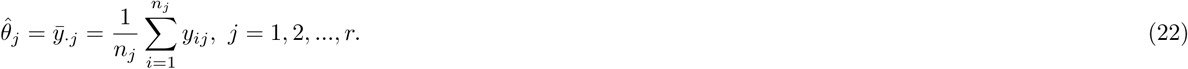

However, in estimating **θ_j_** we do not have to take a dichotomous approach of choosing, based on a preset significance level, between these two extreme choices, the overall weighted mean 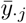 in (21) through full pooling and the separate means 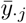 in (21) with no pooling; instead, we could make the assumption that a reasonable estimate to *✓j* lies somewhere along the continuum between 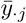 and 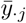, with its exact location derived from the data instead of by imposing an arbitrary threshold. This thinking brings us to the Bayesian methodology.

To simplify the situation, we first assume a known sampling variance *σ^2^* for the ith data point (e.g., trial) for the *j*th entity; or, in Bayesian-style formulation, we build a BML about the distribution of *y_ij_* conditional on **θ_j_**,

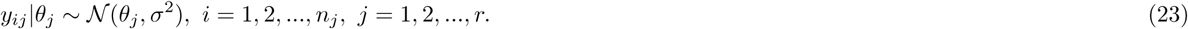

With a prior distribution N*(b_0_,τ^2^)* for **θ_j_** and a noninformative uniform hyperprior for *b_0_* given τ (i.e., *b*_0_|τ ~ 1), the conditional posterior distributions for **θ_j_** can be derived (Gelman et al., 2014) as,

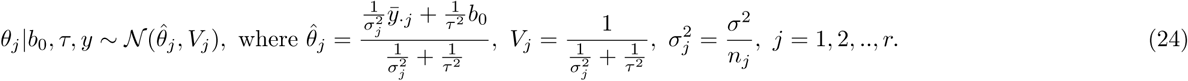

The analytical solution (24) indicates that 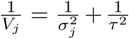, manifesting an intuitive fact that the posterior precision is the cumulative effect of the data precision and the prior precision; that is, the posterior precision is improved by the amount 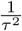 relative to the data precision 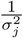. Moreover, the expression for the posterior mode of 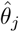 in (24) shows that the estimating choice in the continuum can be expressed as a precision-weighted average between the individual sample means 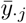 and the overall mean *b*_0_:

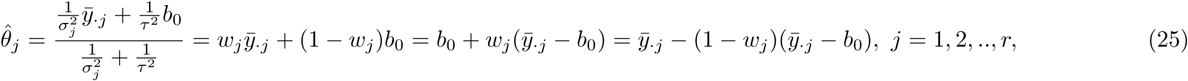

where the weights 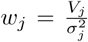 The precision weighting in (25) makes intuitive sense in terms of the previously described limiting cases:

i. The full pooling (21) corresponds to *w_j_* = 0 or τ^2^ = 0, which means that the **θ_j_** are assumed to be the same or fixed at a common value.
ii. The no pooling (22) corresponds to *w_j_ = 1* or τ^2^ = ∞, indicating that the *r* effects **θ_j_** are uniformly distributed within *(—∞,*∞); that is, it corresponds to a noninformative uniform prior on **θ_j_**.
iii. The partial pooling (24) or (25) reflects the fact that the *r* effects **θ_j_** are *a priori* assumed to follow an independent and identically distribution, the prior N*(b_0_, τ^2^).* Under the Bayesian framework, we make statistical inferences about the *r* effects **θ_j_** with a posterior distribution (24) that includes the conventional dichotomous decisions between full pooling (21) and no pooling (22) as two special and extreme cases. Moreover, as expressed in (25), the Bayesian estimate 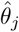 can be conceptualized as the precision-weighted average between the individual estimate 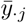 and the overall (or prior) mean *b*_0_, the adjustment of *θ_j_* from the overall mean *b_0_* toward the observed mean 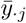, or conversely, the observed mean 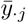 being shrunk toward the overall mean *b*_0_.

An important concept for a Bayesian model is exchangeability. Specifically for the BML (23), the effects **θ_j_** are exchangeable if their joint distribution *p(θ_1_,θ_2_,…,θ_r_)* is immutable or invariant to any random permutation among their indices or orders (e.g., *p(θ_1_,θ_2_,…,θ_r_)* is a symmetric function). Using the ROIs as an example, exchangeability means that, without any *a priori* knowledge about their effects, we can randomly shuffle or relabel them without reducing our knowledge about their effects. In other words, complete ignorance equals exchangeability: before poring over the data, there is no way for us to distinguish the regions from each other. When the exchangeability assumption can be assumed for **θ_j_**, their joint distribution can be expressed as a mixture of independent and identical distributions (Gelman et al., 2014), which is essential in the derivation of the posterior distribution (24) from the prior distribution N*(b_0_,τ^2^)* for **θ_j_**.

To complete the Bayesian inferences for the model (23), we proceed to obtain (i) p(b_0_,τ|y), the marginal posterior distribution of the hyperparameters *(b_0_,τ),* (ii) p(b_0_|τ, y), the posterior distribution of *b*_0_ given τ, and (iii) *p(τ*|y), the posterior distribution of τ with a prior for τ, for example, a noninformative uniform distribution *p(τ)* ~ 1. In practice, the numerical solutions are achieved in a backward order, through Monte Carlo simulations of r to get *p(τ*|y), simulations of *b*_0_ to get p(b_0_|τ, y), and, lastly, simulations of *θ_j_* to get *p(*θ_j_**|b_0_,τ, y) in (24).

### Assessing type S error under BML

In addition to the advantage of information merging across the r entities between the limits of complete and no pooling, a natural question remains: how does BML perform in terms of the conventional type I error as well as type S and type M errors? With the “standard” analysis of r separate models in (18), each effect *θ_j_* is assessed against the sampling variance 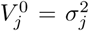. In contrast, under the BML (23), the posterior variance, as shown in (24), is 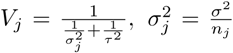 ratio of the two variances 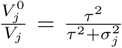 is always less than 1 (except for the limiting cases of σ^2^ → 0 or τ^2^ → ⋡), BML generally assigns a larger uncertainty than the conventional approach with no pooling. That is, the inference for each effect *θ_j_* based on the unified model (23) is more conservative than when the effect is assessed individually through the mode (18). Instead of tightening the overall FPR through some kind of correction for multiplicity among the r separate models, BML addresses the multiplicity issue through precision adjustment or partial pooling under one model with a shrinking or pooling strength of 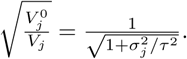

Simulations (Gelman and Tuerlinckx, 2000) indicate that, when making inference based on the 95% quantile interval of the posterior distribution for a single effect *θ_j_* (*j* is fixed, e.g., *j* = 1), the type S error rate for the Bayesian model (23) is less than 0.025 under all circumstances. In contrast, the conventional model (18) would have a substantial type S error rate especially when the sampling variance is large relative to the cross-entities variance (e.g., 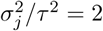); specifically, type S error reaches 10% when 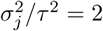, and may go up to 50% if 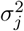 much larger than τ^2^. When multiple comparisons are performed, a similar patterns remains; that is, the type S error rate for the Bayesian model is in general below 2.5%, and is lower than the conventional model with rigorous correction (e.g., Tukey’s honestly significant difference test, wholly significant differences) for multiplicity when σ/τ > 1. The controllability of BML on type S errors is parallel to the usual focus on type I errors under NHST; however, unlike NHST in which the typical I error rate is delibrately controlled through a an FPR threshold, the controllability of type S errors under BML is intrinsically embedded in the modeling mechanism without any explicit imposition.

The model (23) is typically seen in Bayesian statistics textbooks as an intuitive introduction to BML (e.g., Gelman et al., 2014). With the indices i and j coding the task trials and ROIs, respectively, the ANOVA model (19) or its Bayesian counterpart (23) can be utilized to make inferences on an ensemble of ROIs at the individual subject level. The conventional analysis would have to deal with the multiplicity issue because of separate inferences at each ROI (i.e., entity). In contrast, there is only one integrated model (23) that leverages the information among the r entities, and the resulting partial pooling effectively dissolves the multiple testing concern. However, the modeling framework can only be applied for single subject analysis, and it is not suitable at the population level; nevertheless, it serves as an intuitive tool for us to extend to more sophisticated scenarios.

## Appendix E. Derivation of posterior distribution for BML (5)

We start with the BML system (5) with a known sampling variance σ^2^,

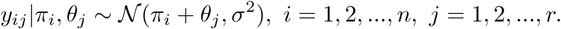

Conditional on *θ_j_* and prior π_i_ ~ N(0,λ^2^), the variance for the sampling mean at the *j*th ROI, 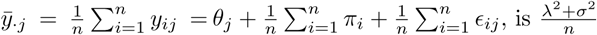 that is,

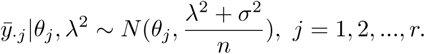

With priors *π_i_* ~ *N(0,λ^2^)* and **θ_j_** ~ *N(μ, τ^2^),* we follow the same derivation as in the likelihood (23), and obtain the posterior distribution,

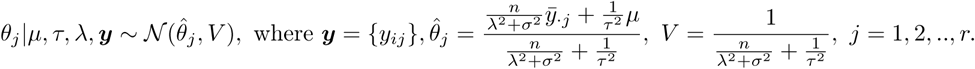

When the sampling variance σ^2^ is unknown, we can solve the LME counterpart in (4),

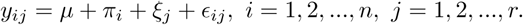

We then plug the estimated variances 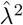, 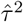 and 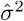 into the above posterior distribution formulas, and obtain the posterior mean and variance through an approximate Bayesian approach.

## Footnotes

1 Entity effects are more popularly called group effects in the Bayesian literature. However, to avoid potential confusions with the neuroimaging terminology in which the word *group* refers to subject categorization (e.g., males vs. females, patients vs. controls) or the analytical step of generalization from individual subjects to the group (corresponding to the word *population* in the Bayesian literature) level, we adopt *entity* to mean each measuring unit such as subject and ROI in the current context.

2 See https://en.wikipedia.org/wiki/Folded-t_and_half-t_distributions for the density *p(v, μ, σ^2^)* of folded non-standardized t-distribution, where the parameters *v, μ, and σ^2^* are the degrees of freedom, mean, and variance.

3 The LKJ prior (Lewandowski, Kurowicka, and Joe, 2009) is a distribution over symmetric positive-definite matrices with the diagonals of 1s.

4 Needless to say, the concept of true effect only makes sense under the current model framework at hand, and may not hold once the model is revised.

5 A single voxel is still possible, but much less likely, to survive this correction approach.

6 A popular cluster reporting method among the neuroimaging software packages is to simply present the investigator only with the icebergsb above the water, the surviving clusters, reinforcing the illusionary either-or dichotomy under NHST.

7 The investigator would not be able to even see such borderline clusters since the typical software implementations mechanically adopt a dichotomous results presentation.

